# Structures of vertebrate R2 retrotransposon complexes during target-primed reverse transcription and after second strand nicking

**DOI:** 10.1101/2024.11.11.623112

**Authors:** Akanksha Thawani, Anthony Rodríguez-Vargas, Briana Van Treeck, Nozhat T Hassan, David L Adelson, Eva Nogales, Kathleen Collins

## Abstract

R2 retrotransposons are model site-specific eukaryotic non-LTR retrotransposons that copy-and-paste into gene loci encoding ribosomal RNAs. Recently we demonstrated that avian A-clade R2 proteins achieve efficient and precise insertion of transgenes into their native safe-harbor loci in human cells. The features of A-clade R2 proteins that support gene insertion are not characterized. Here, we report high resolution cryo-electron microscopy structures of two vertebrate A-clade R2 proteins, avian and testudine, at the initiation of target-primed reverse transcription and one structure after cDNA synthesis and second strand nicking. Using biochemical and cellular assays we discover the basis for high selectivity of template use and unique roles for each of the expanded A-clade zinc-finger domains in nucleic acid recognition. Reverse transcriptase active site architecture is reinforced by an unanticipated insertion motif in vertebrate A-clade R2 proteins. Our work brings first insights to A-clade R2 protein structure during gene insertion and enables further improvement and adaptation of R2-based systems for precise transgene insertion.

## Introduction

Non-long terminal repeat (non-LTR) retrotransposons are mobile genetic elements that are widespread in eukaryotic species. Retrotransposon-derived DNA expression, mobilization, and rearrangement are recognized as major drivers of genome evolution and expansion (*1–3*). In mammals, retrotransposons have expanded via a copy-and-paste mechanism to compose a large portion of genomes. For example, nearly one-third of the human genome originated in the activity of the non-LTR retrotransposon Long Interspersed Element 1 (LINE-1), whose specialized insertion preference for DNA architecture is linked to replication fork progression with a degenerate DNA sequence recognition (*4–6*). The abundant cDNA-derived genome content shapes nuclear organization, chromatin landscape, and transcription of genes and regulatory RNAs (*3*, *7–10*).

Other non-LTR retrotransposons are more target site selective (*11*, *12*). R2 retrotransposons with sequence specificity for insertion to the tandemly repeated ribosomal RNA (rRNA) gene locus (the rDNA) are found within the genomes of multicellular animals including insects, crustaceans and non-mammalian vertebrates (*13*, *14*). R2 protein (R2p) from a moth *Bombyx mori*, hereafter R2Bm, has long been the model system for biochemical characterization of target-primed reverse transcription (TPRT), where nicking of one of the two strands of the target site creates a primer for cDNA synthesis directly into the genome (*15*, *16*) (Fig. 1a). R2p-mediated TPRT was recently re-purposed to insert transgenes into rDNA loci in cultured human cells (*17–20*). This technology, called precise RNA-mediated insertion of transgenes (PRINT), relies on an avian R2p translated from an engineered mRNA co-delivered with a second RNA that templates transgene synthesis (*17*).

**Fig. 1.**
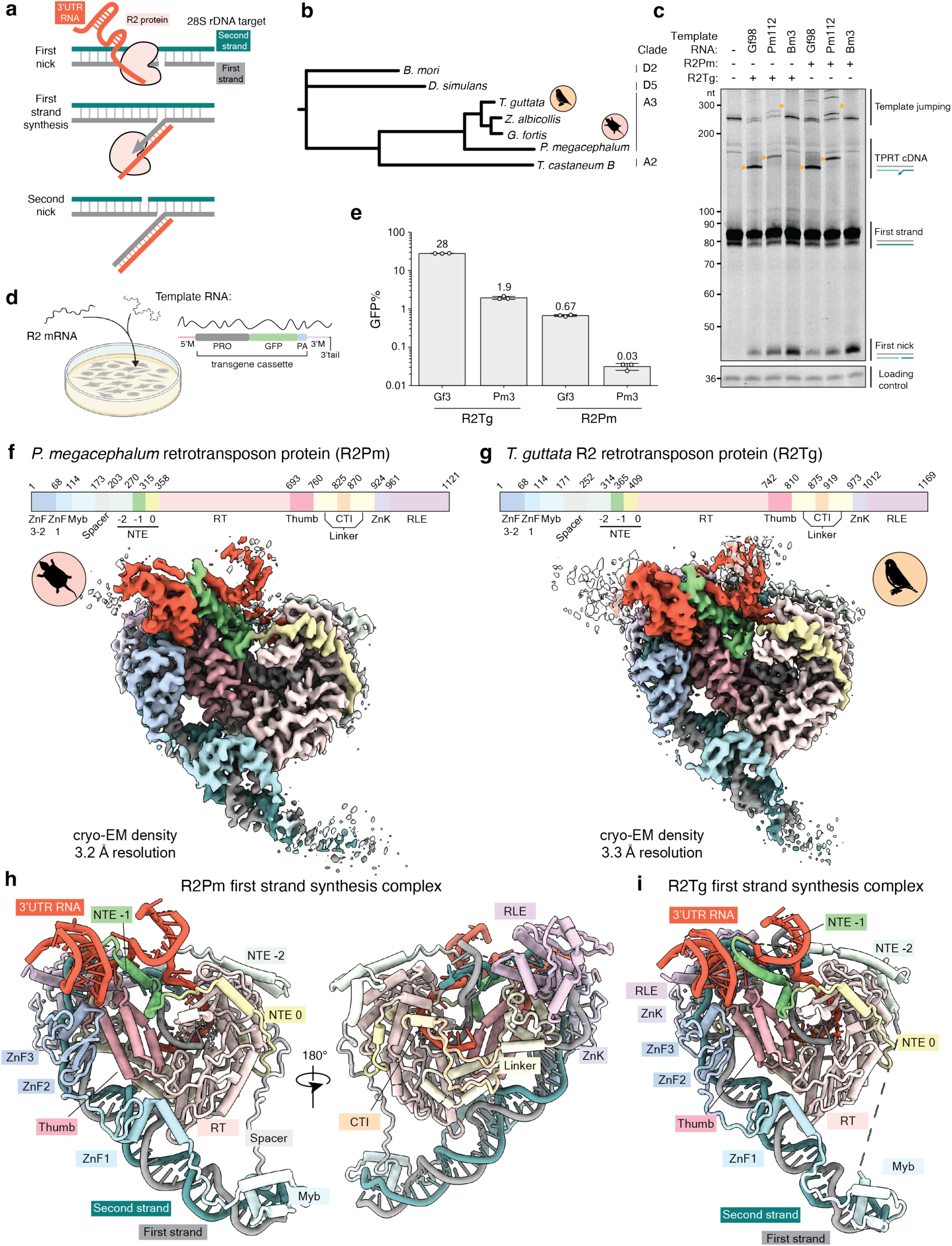
TPRT and PRINT activities and cryo-EM structures of A-clade R2 RNPs initiating TPRT. (a) Schematic of biochemical steps during DNA insertion. (b) Phylogenetic analysis of R2p from the A-clade (birds, turtle, red flour beetle) and D-clade (silk moth and fruit fly) characterized in this and previous work (*17*, *20*). Tree branch length is indicative of substitutions per aligned site. (c) Denaturing PAGE of TPRT reaction products. Orange triangles indicate expected TPRT product lengths for copying a single full-length template (TPRT cDNA). Multiple templates may also be copied in series (template jumping products). R2Pm and R2Tg proteins were assayed with annealed rDNA target site oligonucleotides and different template RNAs, each with an R5 3′ tail: Gf98, Pm112, Bm3. (d) PRINT assay schematic. An mRNA encoding R2Pm or R2Tg protein is transfected with an engineered template RNA comprised of a 5′ module (5′M), modified CMV promoter (PRO), GFP ORF, polyadenylation signal (PA), and 3′ module (3′M) with a 3′tail containing rRNA and A22. (e) PRINT assays with 2-RNA transfection of the R2Pm or R2Tg mRNA and an engineered template RNA with either Gf3 or Pm3 followed by R4A22. Note the log-scale y-axis. (f-g) At top, domains of A-clade R2Pm and R2Tg are illustrated with amino acid numbering; abbreviations given in the text. Cryo-EM density of R2Pm (f) or R2Tg (g) first strand synthesis complex assembled with rDNA target site and either Gf3 full-length 3′UTR RNA (f) or Gf98 RNA (g) is shown and colored by domain. (h-i) Ribbon diagrams of R2Pm (h) or R2Tg (i) first strand synthesis complex structure colored by domains.

The avian R2 retrotransposons belong to the A-clade, which among other clade-distinguishing differences has an expanded number of N-terminal zinc-finger domains (ZnFs) compared to D-clade R2Bm (*13*). Recent structural studies have revealed the architecture of R2Bm ribonucleoprotein (RNP) bound to duplex DNA or launched into TPRT (*21*, *22*), but A-clade R2p remains under-characterized both biochemically and structurally. In particular, the role of the expanded array of ZnFs has not been elucidated, other than its significance for generating a more precise rDNA location of transgene 5′ junction formation with PRINT (*20*). Besides the N-terminal ZnFs, other distinct structural features are likely to exist due to the early divergence of A-clade and D-clade R2s (*13*, *23*). Understanding the structural features and biochemical properties of A-clade vertebrate R2ps will enable rational engineering of these proteins for gene delivery and potential gene therapy applications. Further, while the initial TPRT stage has recently been characterized for R2Bm (*21*), subsequent stages, such as when cDNA synthesis for the first strand has completed and second strand nicking occurs, remain uncharted. In this work, using cryogenic electron microscopy (cryo-EM), we determine structures of A-clade avian (zebrafinch *Taeniopygia guttata*, R2Tg) and testudine (big-headed turtle *Platysternon megacephalum*, R2Pm) R2 RNPs with target site DNA. We also determine R2Pm protein domain configuration after completion of cDNA synthesis and second strand nicking, and we investigate the functional significance of A-clade-specific R2p structural features with biochemical and cellular assays.

## Results

### TPRT and PRINT activities of avian and testudine R2p

R2Tg and also R2p from the white-throated sparrow (*Zonotrichia albicollis*, R2Za) support PRINT (*17*). For comparison to avian R2p, we bioinformatically mined A-clade R2s from reptiles, which are the evolutionary predecessors of Aves (Fig. 1b). We found that testudine and avian R2 3′ untranslated regions (UTRs) have divergent primary sequence but share a possible pseudoknot-hinge-stem loop architecture at the 3′ end of their 3′UTR (Fig. S1a). We assayed the biochemical activities of bacterially expressed and purified R2Tg and R2Pm (Fig. S1b) in combination with 3 RNAs: the optimal avian R2 3′UTR (*17*, *19*) from the medium ground finch *Geospiza fortis* (292 nucleotide (nt) full-length Gf3 or the equally effective Gf98 containing the terminal 98 nt); R2Pm 3′UTR (210 nt full-length Pm3 or shortened Pm112), or R2Bm 3′UTR (248 nt full-length Bm3). Each 3′UTR sequence was followed by 5 nt of downstream rRNA (R5) that can base-pair with primer created by the first strand nick. R2Tg and R2Pm both efficiently used Gf98 and Pm112 RNA for TPRT *in vitro* (Fig. 1c). In competition assays using an RNA mixture for TPRT, both R2p had equal or greater preference for use of Gf98 (Fig. S1c). On the other hand, neither R2Tg nor R2Pm efficiently used Bm3 as a TPRT template (Fig. 1c), suggesting that like R2Tg (*14*), R2Pm has inherent RNA template recognition specificity.

To investigate R2Pm use of template RNA in cells, we tested PRINT efficiency with template RNAs that generate an autonomous GFP expression cassette, comprised of a modified CMV promoter, GFP ORF, and polyadenylation signal. Template RNAs have a 5′ module for RNA stability and a 3′ module with 3′UTR sequence followed by R4 and 22 adenosines (A22) for optimal PRINT (*17*, *19*, *24*). Template RNAs with either Gf3 or Pm3 in the 3′ module were delivered to human RPE-1 cells paired with an mRNA encoding R2Tg or R2Pm (Fig. 1d-e). Template RNA alone gave only background GFP signal (Fig. S1d). R2Tg paired with Gf3 template RNA generated 28% GFP-positive (GFP+) cells, whereas with Pm3 template RNA, only ∼2% of cells were GFP+. R2Pm paired with Gf3 template RNA generated slightly less than 1% GFP+ cells, and as observed for R2Tg, the Pm3 template RNA was used with much lower efficiency (Fig. 1e, Fig. S1d). We conclude that although R2Tg has higher efficiency for transgene insertion than R2Pm, both proteins prefer PRINT template RNA with Gf3. We speculate that this reflects a more favorably homogeneous folding of Gf3 RNA, compared to Pm3 and the previously tested other avian R2 3′ UTRs that all share similar predicted secondary structure (*17*).

### Structures of R2Tg and R2Pm during first strand synthesis

We sought to capture cryo-EM structures of A-clade R2p RNPs during TPRT. We used bacterially expressed and purified R2Pm and R2Tg and Gf3 or Gf98 RNA to assemble TPRT complexes by incubating the proteins with biotinylated rDNA target site duplex (Fig. S2a-b). We halted elongation after 1 nt of cDNA synthesis with dideoxythymidine triphosphate (ddTTP) and isolated complexes using a streptavidin-based pulldown strategy (Fig. S2b). All intended components of ternary complexes were present in the eluted samples, and both proteins had nicked the first strand and initiated cDNA synthesis (Fig. S2c-d). Cryo-EM structure determination for R2Pm with Gf3 in TPRT initiation stage reached an overall resolution of 3.2 Å (Fig. 1f, Fig. S2e, Fig. S3a and Fig. S4a-c). While initial attempts to determine high resolution cryo-EM structure of R2Tg with Gf3 RNA did not succeed due to low particle density, the particle density improved when we use the truncated Gf98 RNA (Fig. S2d, f), and we were able to obtain a structure of R2Tg RNP in the TPRT initiation stage at an overall resolution of 3.3 Å (Fig. 1g, Fig. S3b and Fig. S4a-c). The cryo-EM density maps allowed us to model nearly the entire protein chain for R2Pm and R2Tg as well as the upstream and downstream rDNA and an RNA pseudoknot-hinge-stem fold that forms an extensive surface for protein interaction (Fig. 1h-i, Fig. 2a, Fig. S5a). We also resolved density for ddTTP bound in the active site that is unable to join the cDNA 3′ end due to the incorporated ddTTP (Fig. S5b).

**Fig. 2.**
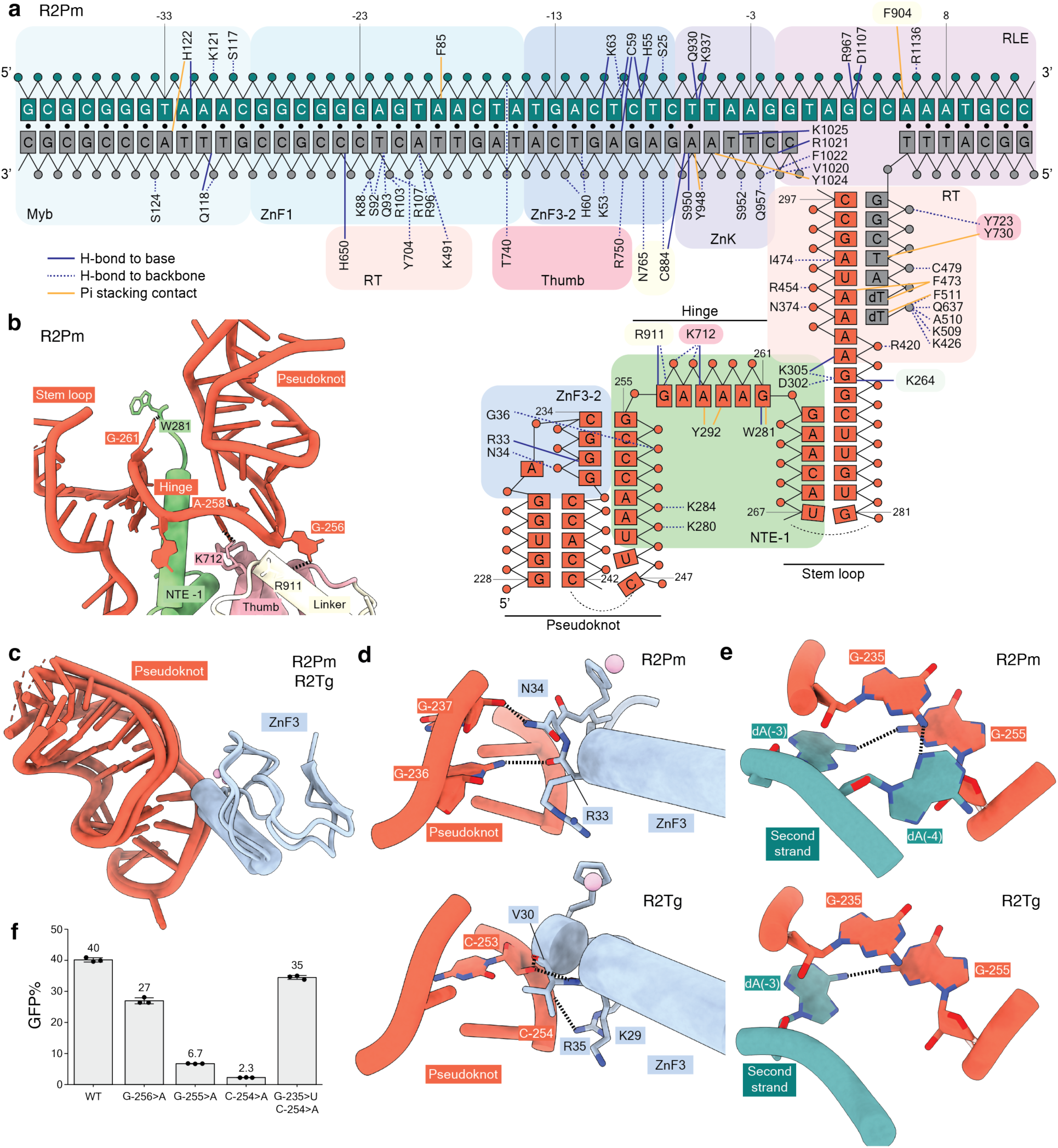
Protein and DNA recognition of R2 3′UTR RNA. (a) Schematic of direct interactions between R2Pm protein, rDNA target site, and 3′UTR RNA in a TPRT initiation complex. Color scheme is consistent with Figure 1. Solid navy lines denote direct hydrogen bonds with the nucleobases or ribonucleobases, while dashed navy lines represent hydrogen bonds with the phosphate backbone or sugars. Solid mustard lines denote pi-stacking contacts with the nucleobases or ribonucleobases. Black circles represent base-pairs in DNA duplex; RNA-DNA or RNA-RNA base-pairing is indicated by apposition. DNA numbering (green and gray strands) is negative upstream or positive downstream of the first strand nick. RNA numbering (red strand) is from the start of Gf3. (b-c) Recognition of the 3′UTR RNA involves the NTE −1, Thumb, Linker and ZnF3 domains. (b) Base-specific hydrogen bonds between bases G-256 and A-258 in the hinge region of 3′UTR RNA and side chains within the Thumb and Linker domains in R2Pm. (c) ZnF3 domain from R2Pm and R2Tg contacts the pseudoknot of 3′UTR RNA. (d) Side chains in ZnF3 make base-specific hydrogen bonds: R2Pm with G-236. R2Pm ZnF3 also makes a contact with the phosphate backbone of base G-237 at the junction of hinge and pseudoknot and R2Tg’s ZnF3 with the phosphate backbone of base C-253. The helix segmentation is an artifact of automated secondary structure assignment. Here and in subsequent figure panels, heteroatom representation has oxygens in red and nitrogens in blue. (e) Base-specific hydrogen bonds between pseudoknot bases and a bases in a single-stranded region of the second strand DNA. (f) PRINT assays using mRNA encoding R2Tg and template RNA with 3′ module Gf98, or a variant Gf98, and R4A22 3′ tail. Base substitutions are numbered according to their position in Gf3, as annotated in (a), with specific mutations described in the main text.

The overall architectures of the A-clade R2p ternary complexes have both similarities and differences with the D-clade R2Bm ternary complex captured at a similar stage of cDNA synthesis (*21*). The shared R2p core domains include the reverse transcriptase (RT) fingers and palm motifs (colored as RT domain) followed by the Thumb, a Linker, the C-terminal zinc-knuckle (ZnK), and the restriction-like endonuclease domain (RLE) (Fig. 1f-i). As shown for R2Bm, the A-clade R2p ZnK and RLE domains melt double-stranded DNA into single-stranded DNA across the first strand nick site. Instead of the two NTEs (NTE 0 and −1) observed in the R2Bm structures (*21*, *22*), The A-clade R2p RT core is preceded by three segments of N-terminal extension (NTE), two previously recognized (NTE 0, −1) and a third (NTE −2) that was not described in the TPRT initiation complex of R2Bm (*21*) or structures of bacterial retroelement proteins (*25*, *26*) (Fig. S5c). NTE motifs are in turn preceded by an evolutionarily variable length of Spacer and the N-terminal ZnF and Myb domains (Fig. 1f-i) that engage rDNA upstream of the first strand nick. Large differences are present, however, in the architecture of A-clade versus D-clade R2p interactions with RNA (see below) and in the shared and unique A-clade R2p ZnF contacts with DNA and RNA that had not been predicted from previous biochemical assays (*20*, *27*). Overall, our structures establish a divergence of A-clade and D-clade R2p nucleic acid interactions.

### RNA recognition by ZnF3 and target site DNA

Of the 292 nt in Gf3 or 98 nt in Gf98, only the region within the 3′ 65 nt is visible in the cryo-EM density map (Fig. 1f, h). The resolved regions of RNA correspond to a 5′ pseudoknot and a 3′ stem connected by a 6 nt hinge (Fig. 2a, Fig. 1f-i). The 4 nt of single-stranded RNA between the 3′ stem and the RNA paired to primer and ddTTP (Fig. S5b) were also resolved. We note that the fold and topology of Gf3 engaged with the two A-clade R2p and of *B. mori* 3′UTR engaged with R2Bm differ significantly, and there is more length of RNA density visible for Gf3 than was visible for RNA bound to R2Bm, potentially due a more stabilized Gf3 3’ end RNA fold (Fig. S5d). The R2p NTE −1, Linker and Thumb domains form a large surface for RNA recognition (Fig. 2a-b, Fig. S5a, e). Key interactions include base-specific hydrogen bonds that Arg911 (R2Pm) or Arg960 (R2Tg) make with G-256, and Lys712 (R2Pm) or Lys763 (R2Tg) make with A-258 in the RNA hinge (Fig. 2a-b, Fig. S5a, e). The sequence specific recognition of GGAAAAG motif in the hinge and adjacent end of the pseudoknot is likely to contribute to the shared template selectivity of avian and testudine R2p.

The A-clade R2p ZnF2 and ZnF3 fold together through a previously unanticipated interaction of beta strands. This folding unit is sandwiched on target site DNA between ZnF1 and RLE (described below) and bookends the RNA pseudoknot from the side opposite NTE −1 (Fig. 1h-i, Fig. 2a-b). ZnF3 contacts the pseudoknot with hydrogen bonding interactions to both backbone and bases (Fig. 2c, d, Fig. S5c). Our structures also reveal that the rDNA target site itself contributes to RNA recognition. We find that bases within the DNA region melted by R2p face toward the pseudoknot. In both R2Pm and R2Tg structures, the base dA(−3) of the second strand creates a sequence-specific hydrogen bond with the base of G-255 at the junction between the pseudoknot and the hinge (Fig. 2e).

To assay the functional significance of the visualized RNA secondary structure and its sequence, we made mutations in the pseudoknot and hinge regions and assessed change in PRINT efficiency. Mutating the hinge base G-256 to A reduced PRINT efficiency and disrupting the pseudoknot base-pairing via G-255 to A or C-254 to A mutation drastically reduced PRINT efficiency (Fig. 2f). Further, restoring the pseudoknot base-pairing with compensatory mutations (G-235>U, C-254>A) restored PRINT activity to a level comparable to the wild-type pseudoknot sequence (Fig. 2f). Altogether, our structural and functional assays demonstrate that multiple regions of the protein recognize and position template RNA, particularly the RNA pseudoknot and the hinge sequence, during the initiation of TPRT.

### Target site recognition by R2 N-terminal DNA binding domains

As also shown for R2Bm in a previous work (*21*), the A-clade R2p ZnK and RLE domains split double-stranded DNA around the first strand nick site (Fig. 3a, Fig. S6a). The nicked first strand upstream of the target site, including its 5′ end, remains buried within the ZnK and RLE domains (Fig. 3a). As a second similarity with R2Bm, the R2Tg and R2Pm motif 6a within the RT domain wedges into a distortion of the upstream target site DNA (Fig. 3b, Fig. 2a, Fig. S5a). Together, the ZnF and Myb domains of A-clade R2p create an extended surface protecting the target site, using the entirety of the 4 domains and also connecting amino acid segments between them (Fig. 3c). In comparison, R2Bm ZnF and Myb domains occupy a much smaller surface of upstream target site, even compared to the A-clade R2p ZnF1 and Myb domains alone (Fig. 3c). A-clade R2p ZnF2 and ZnF3 engage the target site close to the first strand nick site (Fig. 3c, Fig. 2a, Fig. S5a). ZnF2 makes sequence-specific contacts, whereas ZnF1 and ZnF3 predominantly make sequence non-specific contacts with the phosphate backbone of the target DNA (Fig. 3d, Fig. S6b-c). In contrast, R2Bm relies on the ZnF corresponding to the A-clade R2p ZnF1 for sequence-specific contacts (*21*, *22*).

**Fig. 3.**
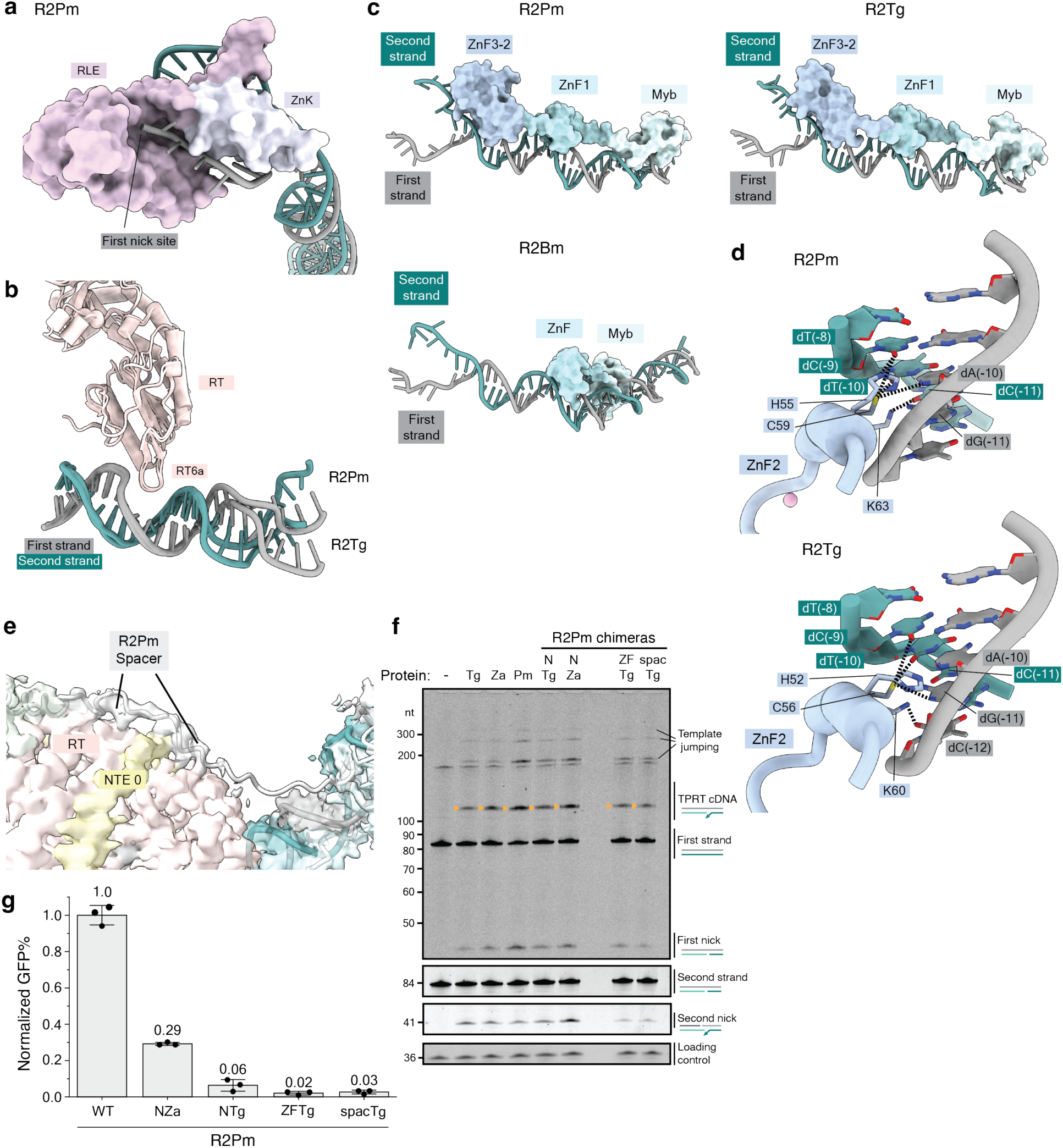
Protein recognition of the target DNA and N-terminal domain requirements for TPRT and PRINT. (a) RLE and ZnK domains surrounding the nicked first strand and single-stranded second strand are illustrated for the R2Pm complex. (b) The motif 6a loop within the RT domain is shown protruding into a distortion in target DNA. (c) Configuration on target DNA of the N-terminal DNA binding domains: the three ZnF and the Myb domain for A-clade R2Pm and R2Tg are compared with the single ZnF and Myb in D-clade R2Bm. (d) Base-reading hydrogen bonds between ZnF2 and the target DNA proximal to the nick site. (e) The unstructured R2Pm Spacer and its interaction with the RT and NTE 0 domains are depicted. (f) Denaturing PAGE of TPRT reaction products with wild-type R2Tg, R2Za, R2Pm and chimeric proteins: R2Pm with the N-terminus (Spacer, Myb, and three ZnFs) from R2Tg (NTg) or R2Za (NZa), R2Pm with ZnF3-2 domains from R2Tg (ZFTg), R2Pm with Spacer from R2Tg (spacTg). Gf68 RNA with R5 3′ tail was used for all assays. Different regions of the same gel are shown, with first strand DNAs and second strand DNAs imaged separately using different 5′ dye. (g) PRINT assays using mRNA encoding R2Pm or the chimeras described in (f). The template RNA 3′ module was Gf3 followed by R4A22.

In previous work using R2Za (*20*), we found that deletion of ZnF2 and ZnF3 had minimal impact on TPRT and reduced, but did not eliminate, PRINT (*20*), suggesting that the ZnF3-2 contacts to RNA and DNA can be lost without severe disruption of RNA and DNA binding specificity. However, removal of ZnF3-2 strikingly decreased the positional precision of transgene 5′ junction formation from the rDNA side (*20*). Contacts between ZnF3 and the pseudoknot would be removed by cDNA synthesis, but ZnF2 contacts to upstream target site could remain (explored below). These contacts, potentially dynamic with continued cDNA synthesis, could influence DNA positioning for second-strand nicking. In concurrence with this idea, a contribution of ZnF3-2 to second strand nicking has been detected using purified proteins under some conditions (*20*). However, future studies are necessary to explore the relationship between R2p’s *in vitro* second strand nicking and productive second strand nicking in cells.

A major difference between the R2Pm and R2Tg structures, in comparison with each other, is the disposition of the Spacer, the region that connects the N-terminal DNA binding domains to the NTE motifs (Fig. 1g-h). R2Tg has a Spacer of ∼80 amino acids that could not be resolved in our cryo-EM map, whereas R2Pm has a Spacer of only ∼30 amino acids that we partially observe in our structure as it makes contacts with the RT core (Fig. 3e). To investigate whether the difference in Spacer length and/or the N-terminal DNA binding domains gives R2Pm its lower PRINT efficiency than R2Tg, we used human cells to express chimeric R2Pm proteins with segments swapped to have an avian R2p Spacer, ZnF3-2, or the entire N-terminal region before the NTE motifs. Purified domain-chimera proteins had similar or slightly better TPRT activity than wild-type R2Pm, but each of the domain-chimera proteins suffered a large loss of PRINT efficiency (Fig. 3f-g). Curiously, R2Pm with the entire N-terminus of R2Tg had substantially lower PRINT efficiency than R2Pm with the entire N-terminus of R2Za, which nonetheless remained compromised for PRINT relative to wild-type R2Pm (Fig. 3g). Altogether, these results demonstrate structural and functional divergence of the N-terminal nucleic acid binding domains and Spacer within vertebrate R2 A-clade proteins to an extent that they are not exchangeable modules of R2p domain architecture. This is suggestive of co-evolution of the catalytic domains with the Spacers and with the DNA binding domains.

### Vertebrate R2p expansion of the C-terminal Insertion

A structural feature specific to the two vertebrate A-clade R2p, relative to R2Bm, is a sequence insertion (hereafter C-terminal insertion, CTI) that threads from after the Thumb to the RT fingers and back (Fig. S7a, Fig. 4a-b). While this Linker sub-region in R2Bm has 11 amino acids connecting two alpha helices, R2Tg and R2Pm have a much longer 44 or 46 amino acids, respectively (Fig. S7a). The CTI anchors to the RT domain with an EWE amino acid triplet (Fig. 4a-b). The R2Pm CTI has a short α-helix that is not present in the R2Tg CTI (Fig. 4a-b). Further, while the entire R2Pm CTI could be easily visualized in the cryo-EM density, the density for the part of the R2Tg CTI that is not facing the RNA-cDNA duplex is only visible at low density thresholds.

**Fig. 4.**
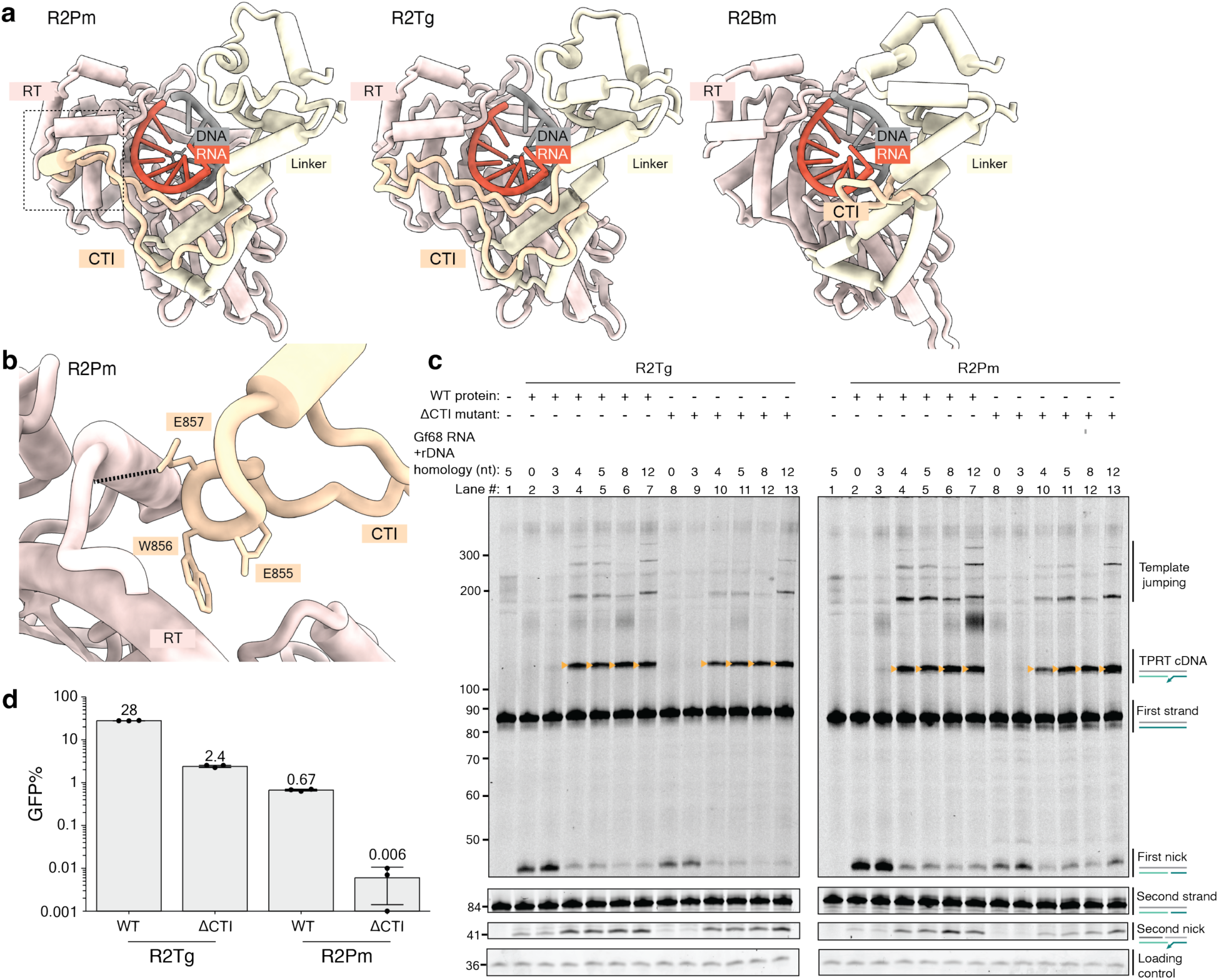
A C-terminal insertion in A-clade R2p. (a) The CTI is rendered in yellow against the RT and Linker domains and RNA:cDNA duplex. The shorter loop present in R2Bm is shown for comparison. (b) Side chains of the conserved EWE motif that anchors the CTI to the RT are displayed for R2Pm. (c) Denaturing PAGE of TPRT reaction products with wild-type R2Tg, R2Tg ΔCTI (CTI truncation) mutant, wild-type R2Pm and R2Pm ΔCTI mutant. Gf68 RNA was synthesized with a variable length of the 3′ tail that base-pairs to target site primer: 0, 3, 4, 5, 8 and 12 nt. Different regions of the same gel are shown, with first strand DNAs and second strand DNAs imaged separately using different 5′ dye. (d) PRINT assays were performed by 2-RNA transfection of the indicated R2p mRNA and template RNA with Gf3 followed by R4A22.

To investigate the functional significance of the longer CTI in A-clade R2p, we truncated the CTI in R2Tg and R2Pm to match the length of this region in R2Bm (ΔCTI mutants) with the goal of deleting the intramolecular EWE anchor without changing the fold of adjacent regions (Fig. S7a). This design was guided by AlphaFold3 (*28*). Wild-type and ΔCTI versions of R2Tg and R2Pm were purified after bacterial expression and assayed for TPRT using Gf68, with the minimized 68 nt of pseudoknot-hinge-stem loop sequence. Due to CTI positioning, we reasoned that it could underlie the previously described avian R2p requirement for base-pairing of primer with the template 3′ tail (*17*). We tested TPRT with Gf68 RNAs harboring different lengths of downstream rRNA (Fig. 4c). In agreement with our previous findings, a 3′ tail of R4 but not R0 or R3 supported TPRT activity of wild-type R2Tg, and the same specificity was observed for wild-type R2Pm (Fig. 4c, lanes 1-3). Additionally increasing the homology length to R5, R8, or R12 had little if any influence on first-strand nicking or cDNA synthesis (Fig. 4c, compare lanes 4-7; note that the adenosine present at the 3′ end of R8 inhibits template jumping). Curiously, CTI truncation did not alter TPRT dependence on R4, but it did decrease unproductive first-strand nicking when the template RNA 3′ tail was too short to support productive TPRT (Fig. 4c, lanes 8-13).

In contrast to reconstituted TPRT, PRINT by both R2Tg and R2Pm was severely inhibited by CTI truncation (Fig. 4d). The percentage of full-length transgene insertions was not proportionally reduced comparing wild-type and ΔCTI R2Tg proteins (Fig. S7b), suggesting that the PRINT deficit is not caused by a substantially lowered processivity of cDNA synthesis in cells. Altogether, our findings lead to the hypothesis that CTI expansion stabilized the active RT fold in a manner critical for PRINT but not limiting for TPRT activity in reactions with purified protein. In a recent study (*18*), the R2Tg CTI was assigned to be a disordered loop and used as a location for insertion of accessory protein modules to optimize transgene insertion. Results from our assays of R2p structure and function above recommend against CTI disruption, which we find to decrease rather than increase transgene insertion efficiency.

### Structure of R2Pm after cDNA synthesis and second strand nicking

Structural insight into a stage of the retrotransposon insertion process following initiation of TPRT is lacking for any clade of R2. We first assayed whether R2Tg and R2Pm proteins had second strand nicking activity dependent on the catalytic activity of the endonuclease domain. Second strand nicking has been demonstrated for R2Bm and recently for two A-clade R2p (R2Za and R2p from flour beetle *Tribolium castaneum*) but is weak compared to first strand nicking (*16*, *20*). We designed a first strand pre-nicked target site DNA with different dye labels at the top and bottom strand 5′ ends. We purified wild-type R2Tg and R2Pm proteins as well as RT or RLE active-site mutants (RTD and END, respectively). When combined with the target site DNA and Gf68 RNA, wild-type and RTD proteins, but not END proteins, nicked the second strand (Fig. 5a). Second strand DNA nicking improved when the wild-type protein was able to perform first strand synthesis upon addition of dNTPs (Fig. 5a). Based on denaturing PAGE migration of the cleavage products, the position of second strand nicking in vitro is similar to the 2 bp offset from the first strand nick detected for all other R2p assayed to date (*16*, *20*)..

**Fig. 5.**
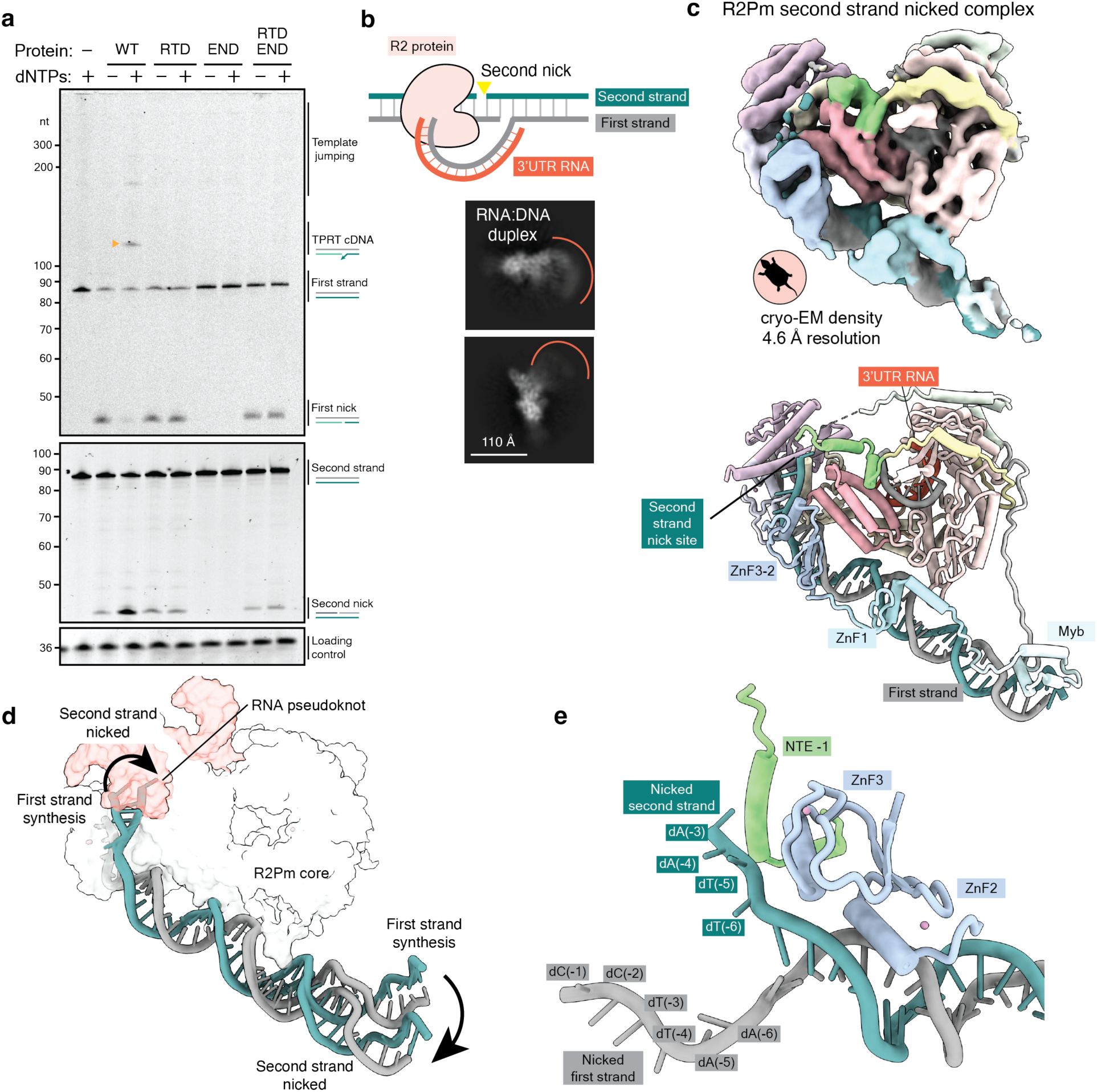
Biochemical activity and cryo-EM structure of A-clade R2 retrotransposon during second strand nicking. (a) Denaturing PAGE of target site nicking and TPRT reaction products from assays using wild-type R2Tg or its RTD and END variants. Gf68 RNA with R5 was used as template. Different regions of the same gel are shown, with first strand DNAs and second strand DNAs imaged separately using different 5′ dyes. Small triangle (mustard) indicates TPRT cDNA. (b) Nucleic acid substrate design to capture a post-TPRT structure for an R2Pm complex. 2D class averages from cryo-EM analysis are shown with inferred range of positions of RNA:cDNA duplex exiting the protein density. (c) Cryo-EM density and ribbon diagram of R2Pm second strand nicked complex assembled, colored by domains. (d) Comparison of upstream target site DNA position in the R2Pm first strand synthesis complex versus second strand nicked complex relative to the R2Pm (NTE to RLE) core (white) and bound 3′UTR RNA (red). After second strand nicking, the nicked single-stranded second strand DNA is displaced towards the RT core and the double-strand DNA bend angle changes near the ZnF1 and Myb domains. (e) Nicked ends of upstream target site DNA are illustrated with nearby R2Pm protein regions NTE −1 and ZnF3-2.

For structure determination, we assembled R2Pm with nucleic acid substrates that mimic the completion of first strand synthesis. The first strand cDNA was annealed to Gf68 with an R5 3′ tail (Fig. 5b, Fig. S8a). The template RNA had a single-nt 5′ overhang that a functional R2p complex would use to complete cDNA synthesis. We added dideoxycytidine (ddCTP) to allow cDNA synthesis completion and then purified complexes and analyzed their composition by denaturing PAGE (Fig. S8b). Some complexes included an intact second strand, but complexes with the second strand nicked were also evident (Fig. S8b). The cryo-EM density reconstructed was for a complex with the second strand nicked. The cryo-EM structure of R2Pm after cDNA synthesis and second strand nicking had an overall resolution of 4.6 Å (Fig. S8c, Fig. S9 and Fig. 5b-c). While some of the 2D class averages visualize the long RNA:cDNA duplex emerging from the protein density, the full length of this duplex was not resolved in 3D reconstructions due to flexibility (Fig. 5b-c).

Our structure revealed a configuration of R2p with the ZnF and Myb domains still bound to upstream target site, as they were at the launch of TPRT. However, considering the entire length of double-stranded DNA, change in its overall positioning was evident comparing R2Pm structures at the start of first strand synthesis and after second strand nicking (Fig. 5d). The single-stranded region of the upstream second strand could be traced towards the first-strand nick site until base dA(−3), consistent with second strand nicking 2 bp upstream from the first strand nick (Fig. 5e). Of note, upon second strand nicking, the 3′ end of the second strand moves into a position occupied by the template RNA pseudoknot at the initiation of first strand synthesis, closer to ZnF3-2 and NTE −1 (Fig. 5d-e). We propose that this positioning would enable R2p to protect the nicked second strand 3′ end from exposure to DNA repair machinery.

## Discussion

### Structural adaptations in R2 evolution

In this study, we investigated the structural basis for steps in the site-specific insertion mechanism of A-clade R2 retrotransposons, which are in a different clade from the best studied D-clade R2Bm system due to an expanded array of N-terminal ZnFs (*12*, *14*). Observations from our cryo-EM along with biochemical and cellular assays demonstrate that each of the three A-clade R2p ZnFs have entirely different nucleic acid recognition principles and roles during gene insertion. We find that these ZnFs, when assayed together in full-length protein context, occupy distinct positions along the upstream rDNA target site. While two ZnFs, together with the Myb domain, bind an extensive length of double-stranded target site DNA, the most N-terminal ZnF, ZnF3, interacts primarily with a newly determined pseudoknot of 3′UTR RNA. Although the ZnF of R2Bm R2p, which corresponds to A-clade ZnF1, has sequence-specific contacts with DNA, it is ZnF2 that has these specific contacts in vertebrate A-clade R2p. The A-clade is believed to be more ancestral than the D-clade (*29*), suggesting that loss of the most N-terminal A-clade ZnFs was accompanied by gain of sequence-specific interaction by the solo D-clade ZnF. Loss of ZnF3-2 may have enabled the D-clade Myb-ZnF DNA binding domains to develop sequence-specific interaction with both downstream and upstream target site sequences (*20*, *30*).

Our work highlights structural differences among the R2p studied at the biochemical level to date, with differences both across clades and also among vertebrate A-clade R2p. Included among these differences is the variable disorder of the Spacer bridging the N-terminal DNA binding domains with the RT-RLE. Unexpectedly, the Spacer and DNA binding domains do not appear to function as a module separable from the RT-RLE. Another particularly divergent structural feature is the CTI. It is of high interest to investigate CTI sequence and structure across a wider diversity of R2p and link this diversity to functional differences at the biochemical and cellular levels.

### R2 retrotransposition and PRINT

Our cryo-EM structure of an R2p complex after second strand nicking reveals that an A-clade R2p remains bound to the upstream target site even after first strand synthesis and second strand nicking. This would ensure close proximity and protection of the upstream and downstream sides of an R2 insertion site during cDNA synthesis. accomplished by a single R2p retained at the site of its initial recruitment. Repositioning of the nicked second strand 3′ end does not place it near the RT active site; instead, the second strand 3′ end is in a protected position that it can occupy after TPRT removes the initially bound pseudoknot-hinge-stem loop RNA. While R2p can make an appropriately positioned second strand nick *in vitro*, questions of whether R2p makes the second strand nick in cells, and if so whether this is mediated by the initially recruited R2p or by a second R2p acting in concert, remain unresolved. As a correlation, deletion of ZnF3-2 inhibits second strand nicking under some conditions *in vitro* and strongly decreases the fidelity of 5’ junction formation for transgenes inserted by PRINT (*20*). However, loss of fidelity in 5’ junction formation could also result from increased R2p dissociation from the upstream target site during cDNA synthesis. Future studies are necessary to explore the mechanisms of second-strand nicking and synthesis in cells.

As a working model, we propose that the persistent upstream binding of A-clade R2p protects otherwise free DNA strand ends but does not launch second strand synthesis. The expanded A-clade R2p ZnF-array recognition and protection of upstream target site DNA could contribute to the favorable function of avian A-clade R2p in transgene insertion by PRINT. However, as A-clade R2Tg and R2Pm have similar RNA binding specificities and similar DNA binding domain configurations on the target site, yet differ strikingly in their ability to support PRINT, factors inherent to the RT-RLE core of R2p are also relevant for efficient PRINT. To develop a site-programmable transgene insertion technology that exploits efficient R2p TPRT in human cells, one possibility would be to replace or supplement the ZnF array with heterologous sequence-specific DNA binding domains, adopting a design principle from zinc-finger nucleases and transcription activator-like effector nucleases (*31*, *32*). Yet, this is unlikely to be straightforward given the lack of domain modularity evident in the deleterious Spacer and DNA binding domain chimeras assayed to date. In combination with extensive target site DNA recognition, the high specificity of vertebrate A-clade R2p for template use by copying the terminal region of 3′UTR RNA would be beneficial for selective insertion of the intended transgene. Our findings inform future improvements and possible reprogramming of R2p-based transgene insertion to the human genome.

## Acknowledgements

We thank members of the Collins and Nogales laboratories for discussions in this collaborative project. We thank Isabella Bartmess for preliminary studies of pseudoknot significance. We thank D. Toso and R. Thakkar at the Cal-Cryo EM facility at UC Berkeley for help with EM data acquisition.

## Author contributions

Conceptualization: A.R.V., A.T., K.C.; Methodology: A.T., A.R.V., B.V.T. and N.T.H.; Investigation: A.T., A.R.V., B.V.T. and N.T.H.; Visualization: A.T., A.R.V., B.V.T. and N.T.H.; Supervision: K.C., E.N. and D.L.A.; Writing—original draft: A.T., A.R.V., and K.C.; Writing— review & editing: all authors.

## Data Availability

The cryo-EM maps reported in this work are deposited under EMD-XXXX, EMD-XXXX and EMD-XXXX in the Electron Microscopy Data Bank and the corresponding atomic model under PDB YYY, PDB YYY and PDB YYY on the Protein Data Bank. All other datasets generated and analyzed during the current study are available from the corresponding authors on request.

## Methods

### Testudine R2 retrotransposon identification

BLASTN+ searches used avian R2 sequences as queries against testudine genome assemblies including *Platysternon megacephalum* (sensitive search, word size = 7) (*33*). Top hits flanked by 28S rRNA were annotated as full-length and the open reading frames were translated using ExPASY (*34*). R2 used for downstream study was selected based on ORF completeness and conservation of essential residues.

### Protein Expression and purification

Construct sequences used in this work are provided in Table S1. Codon-optimized R2 ORFs and other DNA modules were purchased from GenScript. R2 ORFs were cloned into a pET45b vector with N-terminal His14-MBP-bdSUMO tags and C-terminal TwinStrep for bacterial expression (Addgene vector #176534). R2 plasmids were transformed into BL21(DE3) *E. coli* and expressed in modified Terrific Broth media with autoinduction as described previously (*21*). 1L *E. coli* cells were lysed with sonication and the lysate was clarified by centrifugation at 30,000 rpm in Ti45 rotor (Beckman Coulter) for 30 minutes.

For cryo-EM analysis of the R2Tg TPRT initiation and R2Pm second strand nicked complexes, the proteins were purified were purified with the Strep-tactin Superflow Plus resin (Qiagen) and eluted by cleavage with desthiobiotin. For cryo-EM analysis of R2Pm TPRT initiation complex, the protein was purified with NiNTA resin (Qiagen), followed by elution with imidazole. All eluates for cryo-EM analyses were subjected to further purification on a Heparin column (Cytiva) to remove contaminating nucleic acids. Peak elution fractions were analyzed on SDS PAGE, concentrated, flash frozen in liquid and stored in −80°C. Protein concentrations were determined by analyzing with Bradford reagent (Biorad) against a known Bovine Serum Albumin standard.

For *in vitro* TPRT we used predominantly bacterially expressed proteins purified with a single step of Strep-tactin Superflow Plus resin (Qiagen) contained in a gravity-flow column (Bio-rad), which was washed and eluted following the resin manufacturers’ protocol and compatible buffers described previously (*21*). The N-terminal solubility tag was retained for *in vitro* assays since the presence or absence of the tag did not affect TPRT results. The domain-chimera proteins were expressed in and isolated from HEK293T cells as a direct parallel to PRINT assay conditions. N-terminally 1xFLAG-tagged proteins were purified using FLAG antibody resin and determined for concentration as described previously, without modifications (*17*, *35*). Proteins were flash frozen in liquid and stored in −80°C and protein concentrations were determined by densitometry analysis using ImageJ.

The protein mutations made in this study included large truncations (ΔCTI), double alanine substitutions (R2Tg RTD, END) or entire segments swapped between proteins (R2Pm chimeras). For ΔCTI in R2Tg and R2Pm, we truncated positions P884-F914 and P833-Y865, respectively. For R2Pm chimeras, we swapped R2Pm residues Q1-G204 with protein segment M1-Q252 from R2Tg or M1-G242 from R2Za. Additionally, ZnF3-2 motifs within the R2Pm chimeras (Q1-P72) were substituted for a similar region from R2Tg (M1-P70). For the swap of theSpacer region of R2Pm (segment L170-G204), we replaced it with R2Tg protein segment K171-Q252. R2Tg END wasthe combination D1054A, D1067A and RTD was the combination D657A, D658A.

### RNA transcription and purification

Nucleic acid sequences used in this study are provided in Table S1. The 3′UTR sequences of the vertebrate R2 retrotransposons were PCR amplified from parent vectors to include the T7 RNA polymerase promoter. All RNAs were transcribed with T7 RNA polymerase in 40-60 μl reactions with HiScribe T7 High Yield RNA Synthesis Kit (NEB). The *in vitro* transcription reaction was performed for 5 hours at 37°C. The template DNA was removed with DNase RQ1 (Promega), and the transcribed RNA was separated on an 8-12% denaturing polyacrylamide gel. The RNA band was excised and eluted with RNA elution buffer (300 mM NaCl, 10 mM Tris pH8, 0.5% SDS, 5 mM EDTA) overnight at 4°C. The RNA was supplemented with 25 μg glycogen, precipitated with 100% ethanol, centrifuged, and washed with 70% ethanol. The precipitated RNA was air dried before being dissolved in RNase-free H_2_O and if used for cryo-EM supplemented with Ribolock (ThermoFisher) prior to storage at −20°C. Integrity of purified RNA was verified by denaturing PAGE and SYBR Gold nucleic acid gel stain (Thermo Scientific), which was detected by scanning with Typhoon 5 (Cytiva).

### Preparation of TPRT DNA substrates for *in vitro* assays

Oligonucleotide duplex strands (IDT) used in this study have a 3′ block to prevent cDNA synthesis without target-site nicking (Table S1). Target DNA for *in vitro* assays was an 84 bp duplex with both of its strands labeled on their 5′ ends with fluorescent dyes that had non-overlapping emission spectra. For the first strand the sequence is /5IRD800CWN/ ATTCATGCGCGTCACTAATTAGATGACGAGGCATTTGGCTACCTTAAGAGAGTCATAGTTACTCCCGCCGTTTACCCGCGCTTG /3Phos/. The complementary second strand is/5Cy5/CAAGCGCGGGTAAACGGCGGGAGTAACTATGACTCTCTTAAGGTAGCCAAATGCCTCGTCATCTAATTAGTGACGCGCATGAAT /3Phos/. Before annealing, to improve purity and reduce background signal, we size selected and purified from denaturing PAGE each strand following the same approach as for extracting RNA (see RNA transcription and Purification). To anneal these 84 nt strands we first made 10x stocks of expected duplex DNA resuspended in 50 mM KCl and 1 mM MgCl_2_ before heating ssDNA to 95°C for 1 min, then gradually cooled to 25°C over 1 hour using a thermocycler. These annealed substrates were stored at −20°C until use. For all experiments we used a final concentration of 12 nM of the duplex DNA, except for Figure 5a where concentration was reduced to 5 nM to minimize background signal that could obscure product detection.

### TPRT Reactions

*In vitro* TPRT was performed as previously (*17*, *35*), with modifications. TPRT reactions were assembled on ice in a volume of 20 μL with final concentrations of 25 mM Tris-HCl pH 7.5, 150 mM KCl, 5 mM MgCl2, 10 mM DTT, 2% w/v PEG-6000, 5 or 12 nM target DNA duplex, 400 or 50 nM template RNA, 0.5 mM dNTPs, and 30 nM protein (protein added last). For Figure 5a, one (30 nM) or two proteins (15 nM each) were added simultaneously as the last component in the reaction. Reactions were incubated at 30 °C for 15 minutes before heat inactivation at 70 °C for 5 minutes, followed by addition of 2 µL of 10 mg/mL RNase A, incubation for 15 minutes at 55°C, and dilution with 80 μL of stop solution (50 mM Tris-HCl pH 7.5, 20 mM EDTA, 0.2% SDS) spiked with 5-20 ng of a loading control (LC) oligonucleotide (Table S1). Product DNA was purified by phenol-chloroform-isoamyl alcohol (PCI; Thermo Fisher, catalog no. BP17521-100) extraction and ethanol precipitation with 10 μg glycogen as carrier with snap-freezing with liquid nitrogen. Samples were pelleted at ∼18,000 x g for 15-20 minutes at 4°C and pellets washed with 75% (v/v) ethanol, resuspended in 15 μL 0.5x formamide loading dye (95% v/v deionized formamide, 0.025% w/v bromophenol blue, 0.025% w/v xylene cyanol, 5 mM EDTA pH 8.0). Samples were incubated at 95 °C for 3 minutes then placed on ice before loading half of the sample on a denaturing PAGE gel (9% acrylamide/bis 19:1, 7 M urea, 0.6x TBE). Gel scans used a Typhoon 5 (Cytiva) for dual detection of fluorescent dyes on the same gel. Size markers were detected by performing a subsequent gel scan after 6-minute incubation with SYBR Gold stain (Thermo Fisher, catalog no. S11494).

### R2 RT phylogenetic tree, RNA and protein sequence alignments

R2p sequences used in Figures 1b and S7a were collected from previous publications (*13*, *17*, *23*, *27*) excepting the identification of R2Pm described above. For any R2p without a cryo-EM structure, we used AlphaFold3 (28) to predict domain and motif boundaries. We used MAFFT v7.490 (auto model selection) (https://mafft.cbrc.jp/alignment/server/index.html) to align our amino acid sequences of interest. We then used IQTREE v1.6.11 (https://www.hiv.lanl.gov/content/sequence/IQTREE/iqtree.html) for tree reconstruction with 20 maximum likelihood trees and 1000 bootstraps (ModelFinder −m MFP). We used ‘B. Mori’ as the outgroup. The protein alignment in Figure S7a was generated using MAFFT (v7) and the RNA sequence alignment in Figure S1a was performed using Clustal Omega (https://www.ebi.ac.uk/jdispatcher/msa/clustalo).

### Cell culture

RPE-1 cells were grown in DMEM/F12 (Gibco) supplemented with 10% fetal bovine serum (FBS; Seradigm) and 100 μg/mL Primocin (InvivoGen). Cells were cultured at 37 °C under 5% CO_2_. All cells were tested for mycoplasma contamination and human cell lines were validated by short tandem repeat profiling (Promega, catalog no. B9510).

### RNA production for PRINT

Transgene template RNAs and mRNAs for cellular transfection were made using 1 ug of plamid fully linearized with BbsI (NEB) for 4 h at 37 °C and purified with PCR purification kit (QIAGEN, catalog no. 28106) per 20 μL IVT reaction. R2 protein mRNAs expressed C-terminally 3xFLAG tagged protein and were made with AG Clean cap (TriLink, catalog no. N-7113) per the manufacturer’s protocol using UTR sequences from the BioNTech COVID-19 vaccine mRNA (*36*) and an encoded poly-adenosine tail A_30_. mRNAs encoding R2 proteins had 100% uridine substitution with N1-methylpseudouridine. Template RNAs had 100% uridine substitution with pseudouridine. Canonical ribonucleotides were purchased from NEB and uridine analogs were purchased from TriLink. Transcription reactions were incubated at 37 °C for 2 h, followed by addition of 2 μL RNase-free DNase I (Thermo Fisher, catalog no. FEREN0521). Product RNA was purified by desalting with a quick-spin column (Roche, catalog no. 28903408) followed by PCI extraction and precipitation with final concentration of 2.5 M LiCl. After washing twice with 70% ethanol, RNAs were resuspended in 1 mM sodium citrate (pH 6.5). Concentration was determined by NanoDrop and integrity verified by denaturing urea-PAGE with direct staining using SYBR Gold (Thermo Fisher, catalog no. S11494).

### PRINT by 2-RNA delivery

RPE-1 cells at 50% confluency, in log-phase growth, were replated at 350,000 cells per well in twelve-well plates. Cells were reverse-transfected with mRNA and template RNA using Lipofectamine MessengerMAX at ½ mass/volume ratio as per the manufacturer’s instructions. 0.5 μg total RNA mixture was transfected per well of a twelve-well plate and mRNA/template molar ratio was 1:3. Cells were collected 20-24 hours (1 day) after transfection. Plasmid sequences for mRNA and template RNA transcription are provided in Table S1.

### Flow cytometry

Cells were trypsinized, and trypsin was inactivated by addition of dPBS (-Mg^2+^, -Ca^2+^) supplemented with 0.5 mM EDTA and 2% FBS. Cell samples were then analyzed by Attune NxT Flow Cytometer (Thermo Fisher) under the voltage setting of FSC 70V, SSC 280V, BL1 250V. Data analysis was performed in FlowJo (v. 10.8.1). Cells transfected with template RNA only were used as negative controls. The %GFP+ was calculated by subtracting template-alone %GFP+.

### Genomic (g) DNA purification and ddPCR

Frozen cell pellets were thawed on ice and resuspended in 200 μL of RIPA lysis buffer (150 mM NaCl, 50 mM Tris-HCl pH 7.5, 1 mM EDTA, 1% Tx-100, 0.5% sodium deoxycholate, 0.1% SDS, 1 mM DTT). Each 200 μL of lysate was treated with 10 μL of 10 mg/mL RNaseA (Thermo Fisher, catalog no. FEREN0531) at 37 °C for 30-60 min, followed by incubation with 5 μL of 20 mg/mL Proteinase K (Thermo Fisher, catalog no. FEREO0491) at 50 °C overnight. gDNA was then isolated by extraction with PCI and ethanol precipitation. After centrifugation, the aqueous layer was transferred to a fresh tube containing 50 μg glycogen, to which 1/10 volume 5 M NaCl and 3 vol 100% ethanol were added. gDNA was precipitated at −20 °C for at least 30 min. After a 30 min spin, gDNA pellets were washed 3 times with 70% ethanol, air-dried, and resuspended in TE (10 mM Tris-HCl pH 8.0, 1 mM EDTA).

gDNA was digested for 2 h with BamHI and XmnI (NEB). Multiplex 24 μL ddPCR reactions were prepared by mixing 12 μL of ddPCR supermix (no dUTP; Bio-Rad, catalog no. 1863024), forward and reverse primers for target and reference genes (IDT, 833 nM final concentration each), probes complementary to target and reference amplicons (IDT; FAM for target and HEX for reference, 250 nM final concentration each) and digested gDNA at 1-5 ng/μL final concentration. Oligonucleotide sequences are listed in Table S1. Reaction mix was transferred to a DG8 cartridge (Bio-Rad, catalog no. 1864007) along with 70 uL of droplet generation oil (Bio-Rad, catalog no. 1863005), and droplets were generated in a Bio-Rad QX200 Droplet Generator. Following droplet generation, 40 μL was transferred into a 96-well plate and heat-sealed with pierceable foil. The droplets were thermal-cycled under the manufacturer’s recommended conditions with an annealing and/or extension temperature of 56 °C and analyzed using QX Manager software with default settings. *RPP30* (copy number of 3 in RPE cells) was used as the reference gene for all copy number analysis.

### Pulldown of first strand synthesis complex for cryo-EM analysis

The 76-bp 28S DNA target with 5′ biotinylated second strand was annealed separately. First strand synthesis complex was assembled by incubating 160 nM of pre-annealed 76-bp 28S DNA target, 250-300 nM of R2 protein, 300 nM of 3′UTR RNA, 1 μg/mL bdSumo protease and 100 μM of 2′,3′-dideoxythymidine (ddTTP) in 1ml total volume in pulldown buffer (25 mM HEPES-KOH pH 7.9, 400 mM potassium acetate, 10 mM magnesium acetate, 1 mM TCEP). The complex was assembled on a rotator and incubated for 30 minutes at 37 °C. 80 μl of Streptavidin Sepharose High Performance resin (Cytiva) was pre-washed and incubated with the pulldown reaction at room temperature for 30 minutes. The flowthrough was removed, and the beads were washed twice with 0.5 mL pulldown buffer. The elution was performed for 30 minutes at 37 °C in the presence of 5mM desthiobiotin and 4-5 μL PvuII enzyme. The input, flowthrough, washes and elution samples were analyzed on an SDS PAGE and denaturing PAGE gels and stained with Coomassie blue and SYBR Gold (ThermoFisher) stains, respectively. The pulldown eluate was concentrated to 25-40 μL for cryo-EM grid preparation.

### Pulldown of second strand cleavage complex for cryo-EM analysis

28S DNA target with pre-nicked first strand to mimic synthesized Gf68 cDNA, 5′ biotinylated second strand and Gf68-R5 RNA were annealed separately. Sub-stoichiometric RNA concentration of 0.7x was used to anneal the cDNA substrate. Second strand synthesis complex was assembled by incubating 160 nM of cDNA substrate, 250-300 nM of R2 protein, 1 μg/mL bdSumo protease and 100 μM of 2′,3′-dideoxycytidine (ddCTP) in 1 mL total volume in pulldown buffer (25 mM HEPES-KOH pH 7.9, 400 mM potassium acetate, 10 mM magnesium acetate, 1 mM TCEP). The complex was assembled on a rotator and incubated for 30 minutes at 37 °C. 80 μl of Streptavidin Sepharose High Performance resin (Cytiva) was pre-washed and incubated with the pulldown reaction at room temperature for 30 minutes. The flowthrough was removed, and the beads were washed twice with 0.5 mL pulldown buffer. The elution was performed for 30 minutes at 37 °C in the presence of 5 mM desthiobiotin and 4-5 μL PvuII enzyme. The input, flowthrough, washes and elution samples were analyzed on an SDS PAGE and denaturing PAGE gels and stained with Coomassie blue and SYBR Gold (ThermoFisher) stains, respectively. The pulldown eluate was concentrated to 25-40 μL for cryo-EM grid preparation.

### Cryo-EM grid preparation and data collection

Preparation of graphene oxide grids was adapted from our previously developed protocol (*37*). Briefly, Quantifoil Au/Cu R1.2/1.3 grids 200-mesh (Quantifoil, Micro Tools GmbH, Germany) were cleaned by applying two drops of chloroform, then glow discharged. 4 μL of 1mg/mL polyethylenimine HCl MAX Linear Mw 40k (PEI, Polysciences) in 25 mM K-HEPES pH 7.5 was applied to the grids, incubated for 2 minutes, blotted away, washed twice with H_2_O, and dried for 15 minutes on Whatman paper. Graphene oxide (Sigma, 763705) was diluted to 0.2 mg/mL in H_2_O, vortexed for 30 seconds, and precipitated at 1,200 xg for 60 s. 4 μL of supernatant was applied to the PEI treated grids, incubated for 2 minutes, blotted away, washed twice with 4 μL H_2_O each, and dried for 15 minutes on Whatman paper before using for grid preparation. 4 μL of R2 complex was applied to the freshly prepared graphene oxide coated grid and incubated for 60 s at 12 °C and 100% humidity in a Vitrobot Mark IV (ThermoFisher). The grid was then blotted for 1 s with a blot force of 1 and vitrified by plunging into liquid ethane.

For the R2Pm TPRT initiation complex, micrographs were collected on a Titan Krios microscope (ThermoFisher) operated at 300 keV and equipped with a K3 Summit direct electron detector (Gatan). 6,425 movies were recorded using the program SerialEM at a nominal magnification of 105,000x in super-resolution mode (super-resolution pixel size of 0.405 Å/pixel) and with a defocus range of −1.5 μm to −2.5 μm. The electron exposure was about 50 e^-^/Å^2^. Each movie stack contained 50 frames. The same procedure was followed for the R2Tg TPRT initiation complex to record 5,096 movies. For the R2Pm second strand nicked complex, micrographs were collected on a Talos Arctica microscope (ThermoFisher) operated at 200 keV and equipped with a K3 Summit direct electron detector (Gatan). 9,192 movies were recorded using the program SerialEM at a nominal magnification of 36,000x in super-resolution mode (super-resolution pixel size of 0.57 Å/pixel) and with a defocus range of −1.5 μm to −2.5 μm. The electron exposure was about 50 e^-^/Å^2^. Each movie stack contained 50 frames.

### Cryo-EM Data Processing

Cryo-EM data processing workflows are outlined in Supplementary Figs. 3, 4 and 7. All movie frames were motion corrected using MotionCor2 (*38*) in RELION 3.1.1 (*39*) and the corresponding super-resolution pixels size was binned 2x during this process. Contrast transfer function (CTF) parameters for each micrograph were estimated using CTFFIND4.1 (*40*). Motion corrected micrographs were imported into cryoSPARC v.4.5 and particles were picked using Blob Picker. 2D classification was performed in cryoSPARC. 400,309 particles for the R2Pm first strand synthesis complex, 763,427 particles for the R2Tg first strand synthesis complex, and 77,001 particles for the R2Pm second strand cleavage complex were imported back to RELION, 3D initial models were generated, and 3D classification with alignment was performed for each dataset. The class for the R2Pm second strand cleavage complex with 32,239 particles was further refined. Due to the limited number of particles, no further processing was carried out. For R2Pm and R2Tg first strand synthesis complexes, the classes with the best features were selected, refined, particles were polished with Bayesian polishing, and these classes were subjected to one round of 3D classification without alignment on the entire complex. The best class with sharpest features was selected and refined. The final reconstruction was obtained at 3.2 Å nominal resolution from 30,692 particles for the R2Pm complex, and 3.3 Å nominal resolution from 18,892 particles for the R2Tg complex. The cryo-EM maps were sharpened with post-processing in RELION for model building and display in the figures.

### Model Building and Refinement

Model building was initiated by rigid-body fitting the AlphaFold3 (*28*) model of R2Pm and R2Tg proteins engaged with rDNA target into the final cryo-EM density maps using UCSF ChimeraX (*41*). The R2Pm and R2Tg proteins were first manually inspected in COOT (*42*) and then subjected to real space refinement in PHENIX (*43*). Amino acid side chains were manually inspected in COOT and modified when needed before another round of real space refinement in PHENIX. Ribosomal DNA target and 3′UTR RNA were built starting with the R2Bm structure (PDB 8GH6). The parts of DNA target, particularly the single-stranded DNA, that did not fit the density were built de novo in COOT. RNA sequence was corrected to reflect the sequence used in experimental structures. Parts of the RNA were manually built de novo in COOT. The model was corrected to include an unincorporated dTTP obtained from PDB 1CR1. Both were docked into the density map using UCSF Chimera and manually rebuilt with the corresponding DNA chain in COOT. Four zinc atoms were manually placed in each structure and refined in COOT. The model was subjected to global refinement using iterative rounds of real-space refinements in PHENIX with rotamer and Ramachandran restraints. The complete model was subjected to a final real-space refinement and validation in PHENIX. Model building and validation statistics are listed in Table S2.

### Comparison with *Bombyx mori* R2 RT

*Bombyx mori* R2 RT (PDB 8GH6) was aligned with the vertebrate R2 protein chains using the MatchMaker tool in UCSF ChimeraX.

---

**Fig. S1.**
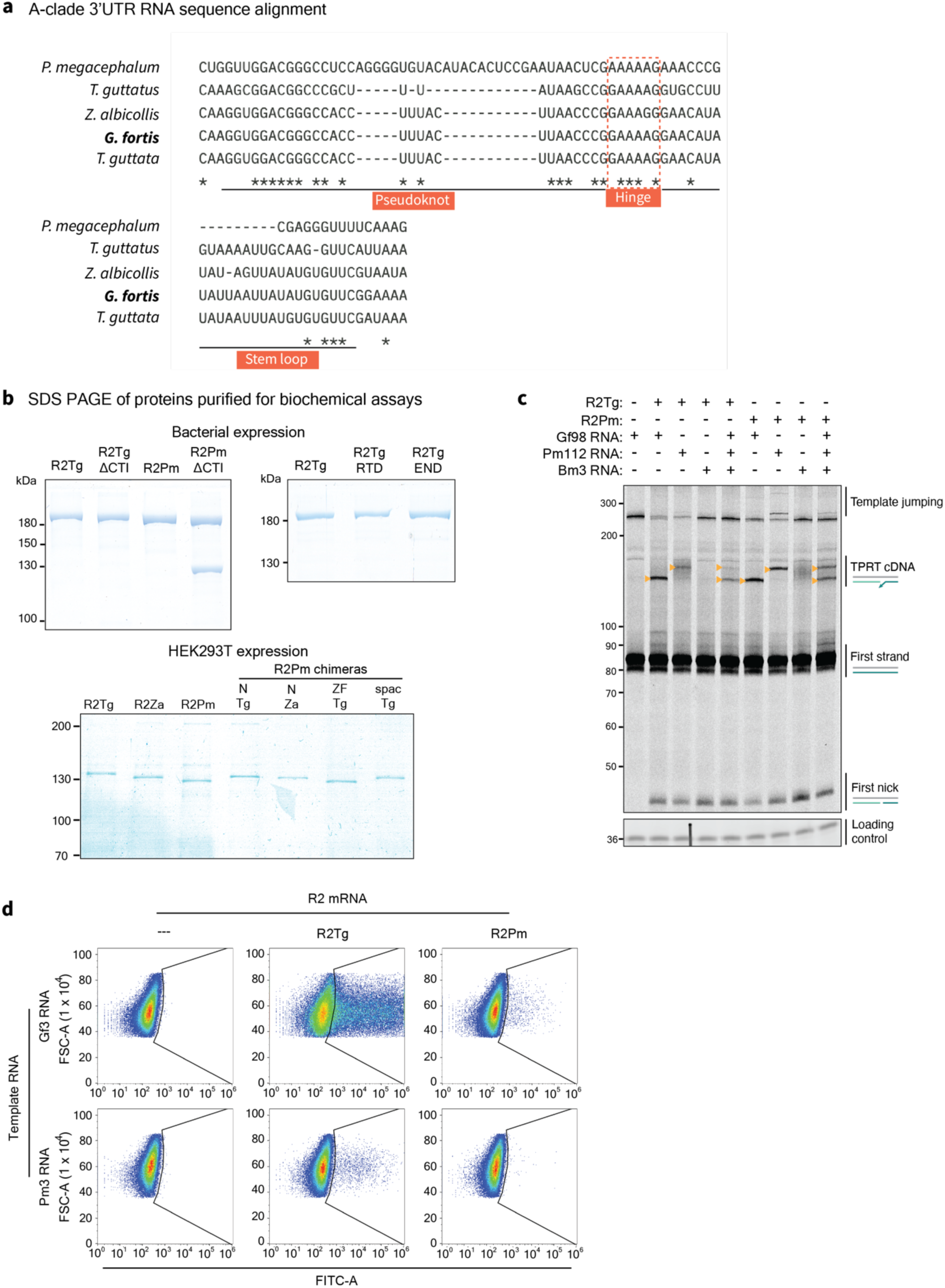
R2 terminal 3′UTR sequence alignment and biochemical assays. (a) Multiple sequence alignment of the 3′-terminal regions of 3′UTR RNAs from A-clade avian (bottom four species) and testudine (*P. megacephalum)* R2, using species with R2p described in the main text or in a previous work (ref: 17). Numbering is from the start of the aligned region only. Nucleotide identity is indicated with an asterisk, and regions of pseudoknot, hinge, and 3’ stem-loop are indicated. (b) Coomassie blue stained SDS PAGE gels showing all wild-type and variant versions of R2p used for TPRT assays. All proteins used for TPRT retained their tag fusions (see Methods). The smaller protein in the R2Pm ΔCTI sample likely reflects increased proteolysis. Purification used the C-terminal Twin-Strep tag, such that an ∼120 kDa protein fragment would lack ZnF3-2 and the Hisx16-MBP-bdSUMO tag of the intact protein; only the full-length protein was quantified to normalize protein concentration. (c) Denaturing PAGE analysis of TPRT reaction products using single or mixed 3′UTR-derived RNA. Gf98, Pm112, and Bm3 are described in the main text, each used here with an R5 3′ tail. Small triangles (mustard) indicate expected TPRT product length for nick-primed cDNA synthesis using a single full-length RNA. Template jumping indicates products from the processive use of additional template(s). The first lane is a mock reaction showing the migration of target site and loading control DNAs; the background bands are not cDNA products. (d) Representative flow cytometry results from one replicate of the Figure 1 PRINT experiment. The gating of GFP+ cells is demarcated with black lines. The x-axis is GFP intensity, and the y-axis approximates cell size. Panels on the far-left show results for cells transfected with template RNA only, without mRNA, as negative controls.

**Fig. S2.**
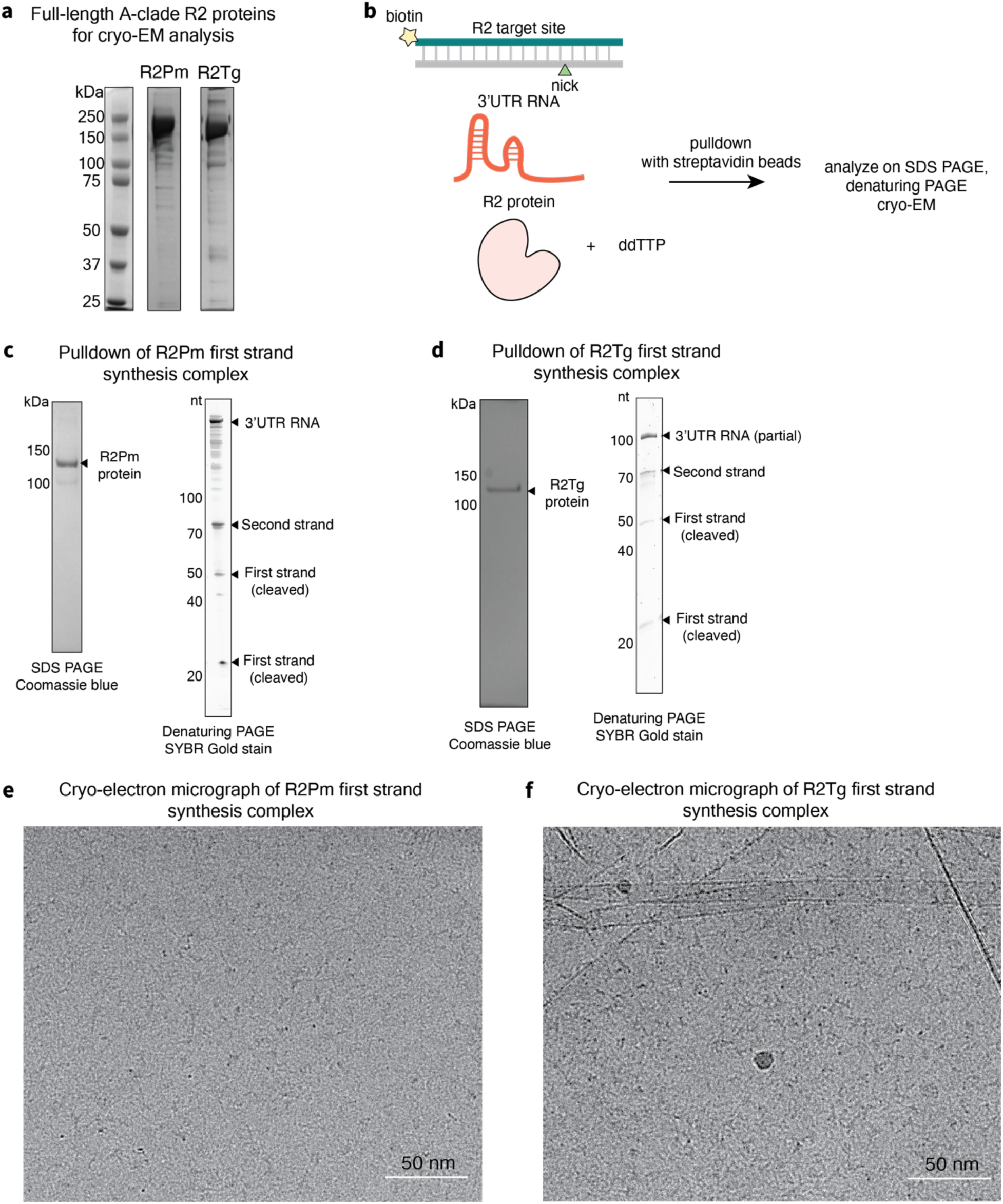
Assembly of TPRT initiation complexes for cryo-EM analysis. (a) SDS PAGE of purified full-length R2Pm and R2Tg proteins after Strep-affinity and Heparin purification for cryo-EM analysis. (b) Schematic of R2 complex assembly during TPRT. R2 proteins were incubated with biotinylated target site DNA, 3′UTR RNA (full-length or truncated) and ddTTP for production of the TPRT initiation state. (c) SDS PAGE analysis of protein and denaturing PAGE analysis of nucleic acids in the pulldown eluate for the R2Pm TPRT initiation complex. Gf3 RNA was used. (e) SDS PAGE analysis of protein and denaturing PAGE analysis of nucleic acids in the pulldown eluate for the R2Tg TPRT initiation complex. Gf98 RNA was used. (e) Representative cryo-EM micrograph of the pulldown eluate for R2Pm captured during TPRT initiation. (f) Representative cryo-EM micrograph of the pulldown eluate for R2Tg captured during TPRT initiation.

**Fig. S3.**
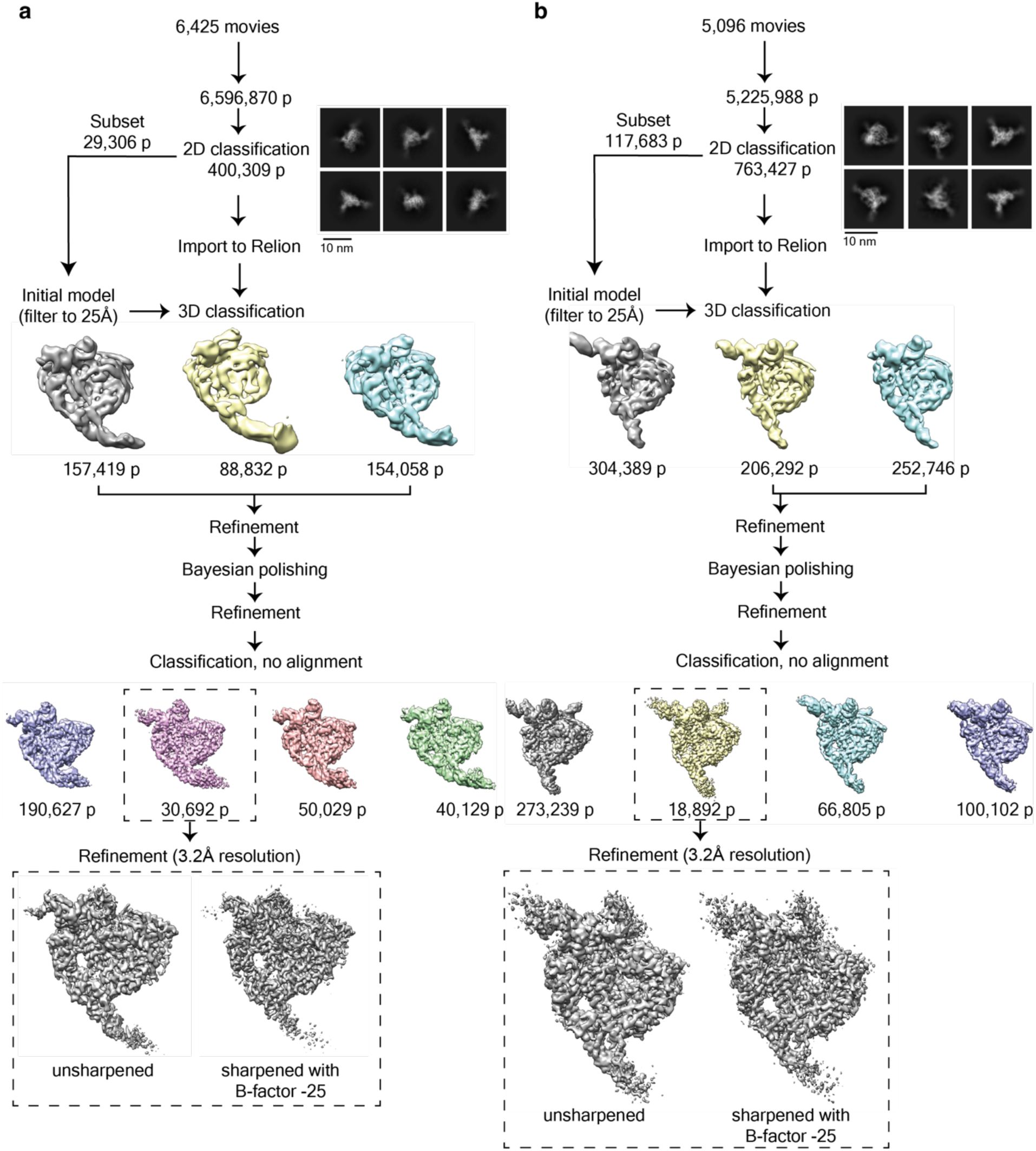
Cryo-EM data processing pipeline used for the R2Pm and R2Tg first strand synthesis complexes. Single particle analysis workflow leading to the reconstruction of the (a) R2Pm and (b) R2Tg first strand synthesis complexes described in Figures 1-4. Densities for the final structures are shown both before and after sharpening.

**Fig. S4.**
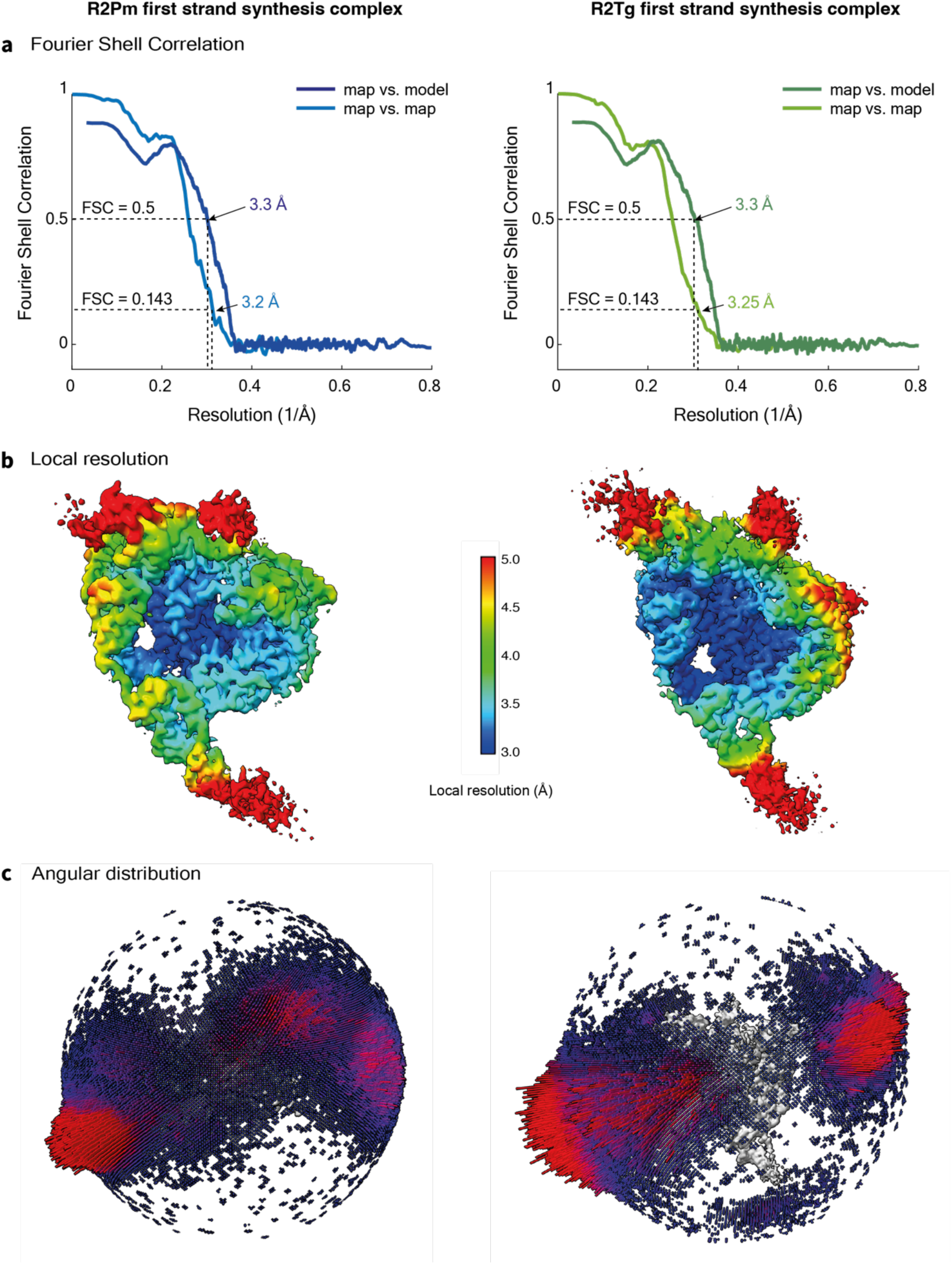
Resolution estimation. (a) Gold-standard FSC curve and map versus model FSC obtained from the final model after validation in Phenix for the R2Pm (left) and R2Tg (right) TPRT initiation complexes. (b) Unsharpened density maps obtained from analysis in Supplementary Figure 3 were colored by local resolution as estimated using Relion 3.1. (c) Particle orientation distribution in the final reconstructions.

**Fig. S5.**
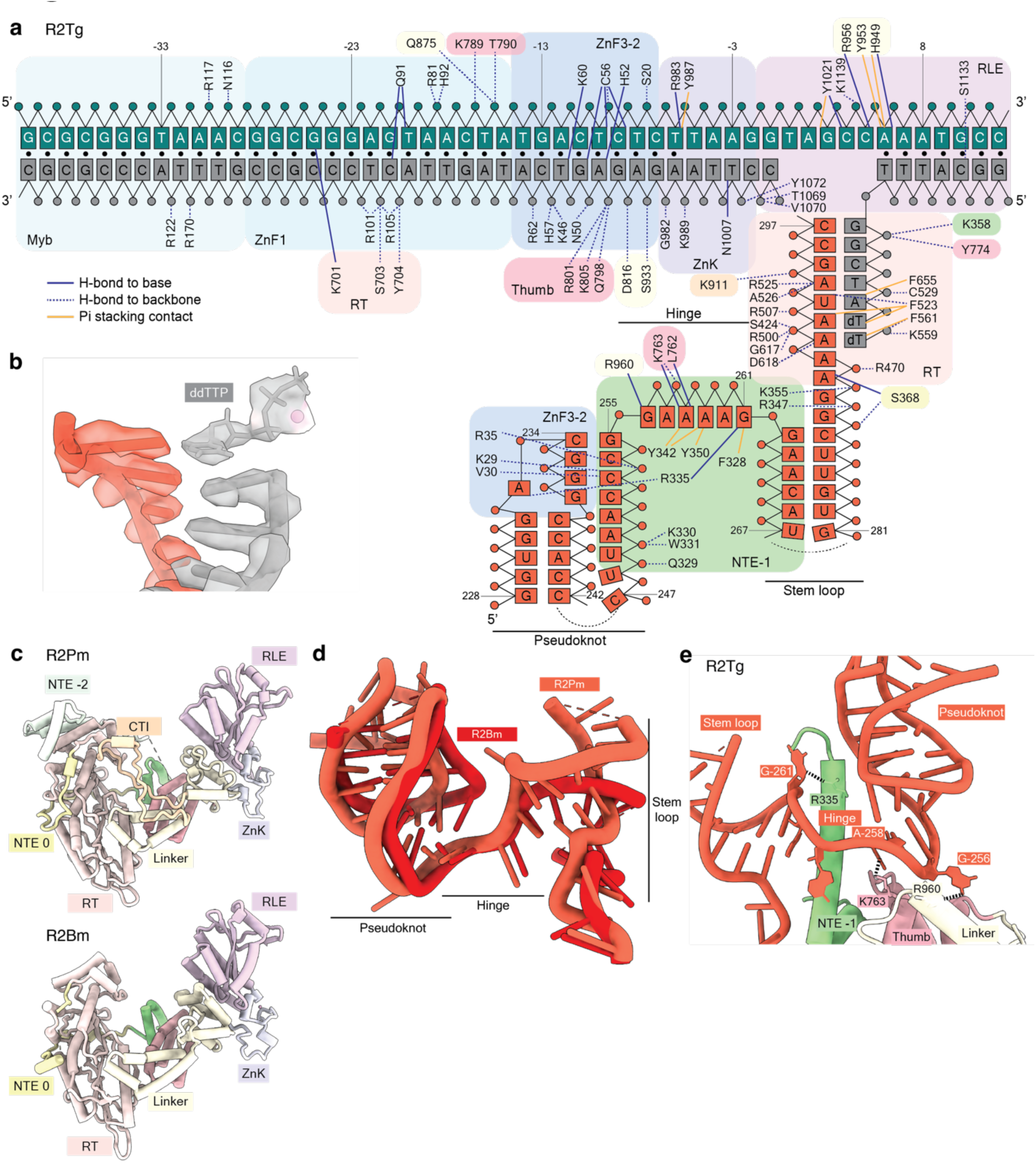
Nucleic acid interactions by the R2Tg protein during TPRT initiation. (a) Schematic of direct interactions between R2Tg protein, target site DNA, and 3′UTR RNA. Color scheme and labeling are consistent with Figure 1. Solid navy lines denote direct hydrogen bonds with the nucleobases or ribonucleobases, while dashed navy lines represent hydrogen bonds with the phosphate backbone or sugars. Solid mustard lines denote pi-stacking contacts with the nucleobases or ribonucleobases. Black circles represent canonically base-paired DNA bases. (b) The RT active site harbored an unincorporated ddTTP that was resolved with a coordinated Mg^2+^ ion (sphere). (c) The RT-RLE core is compared for R2Pm and R2Bm using the region from NTE to C-terminus. Compared to the D-clade R2Bm, A-clade R2Pm contains expanded domains including NTE −2 and CTI. (d) Overlay of RNA backbones and base orientation comparing TPRT initiation complexes of R2Pm (orange-red) and R2Bm (darker red) from PDB 8gh6. The entire protein chain was superimposed. (e) Recognition of 3′UTR RNA in the R2Tg TPRT initiation complex by NTE −1, Thumb and Linker. Base-specific hydrogen bonds occur between bases G-256 and A-258 of the hinge region and side chains within the Thumb and Linker. Compare to R2Pm in Figure 2b.

**Fig. S6.**
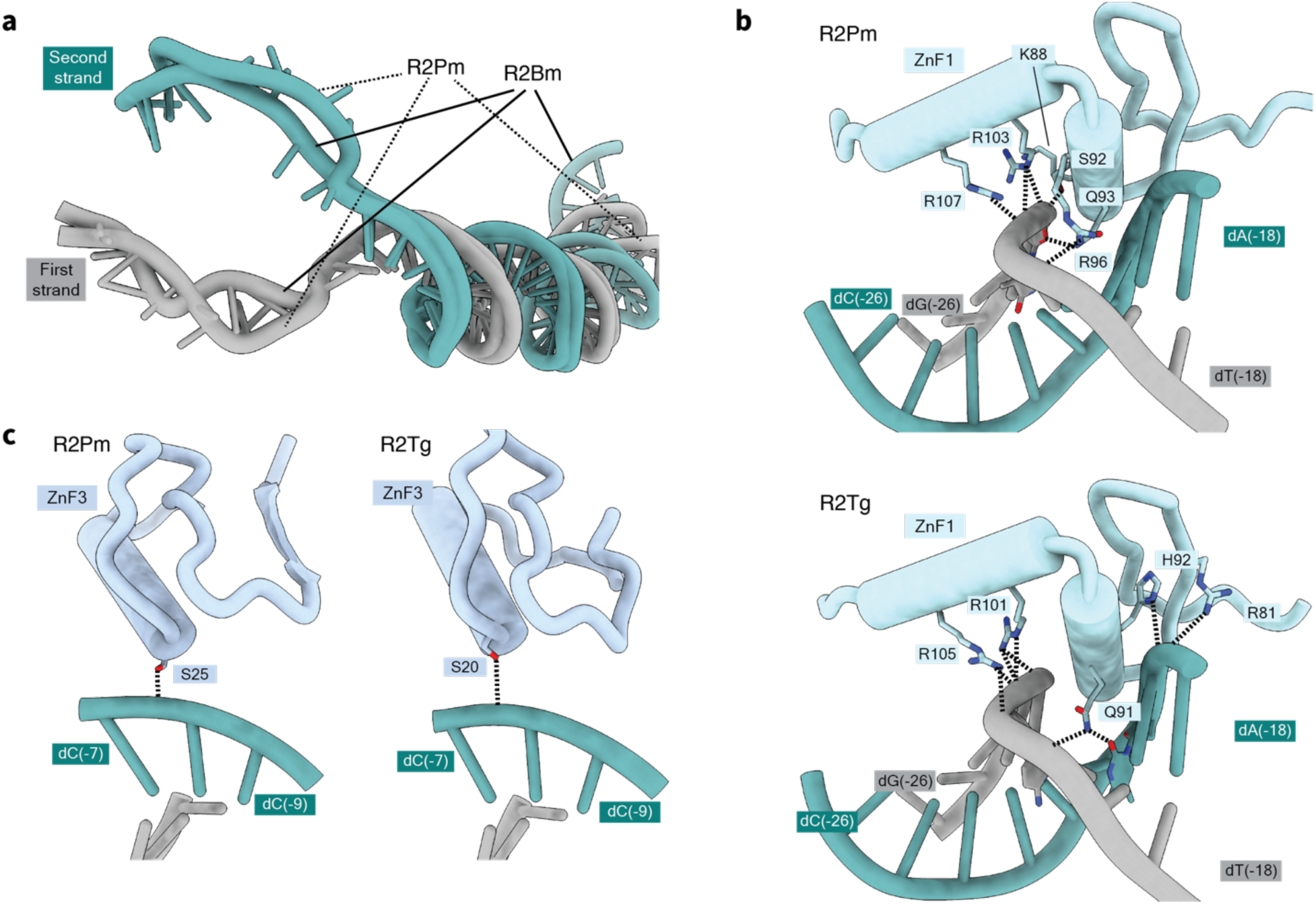
Target DNA engagement by R2 proteins. (a) DNA upstream from the first nick site in the R2Pm TPRT initiation complex was superimposed with the equivalent upstream DNA in the R2Bm TPRT initiation complex (PDB 8gh6) for comparison. (b) Target site recognition by ZnF1 occurs predominantly by sequence non-specific hydrogen bonds with the DNA backbone, shown R2Pm at top and for R2Tg below. R2Tg Q91 side chain makes one base-specific contact. (c) ZnF3 has minimal, sequence non-specific hydrogen bonds with the DNA backbone.

**Fig. S7.**
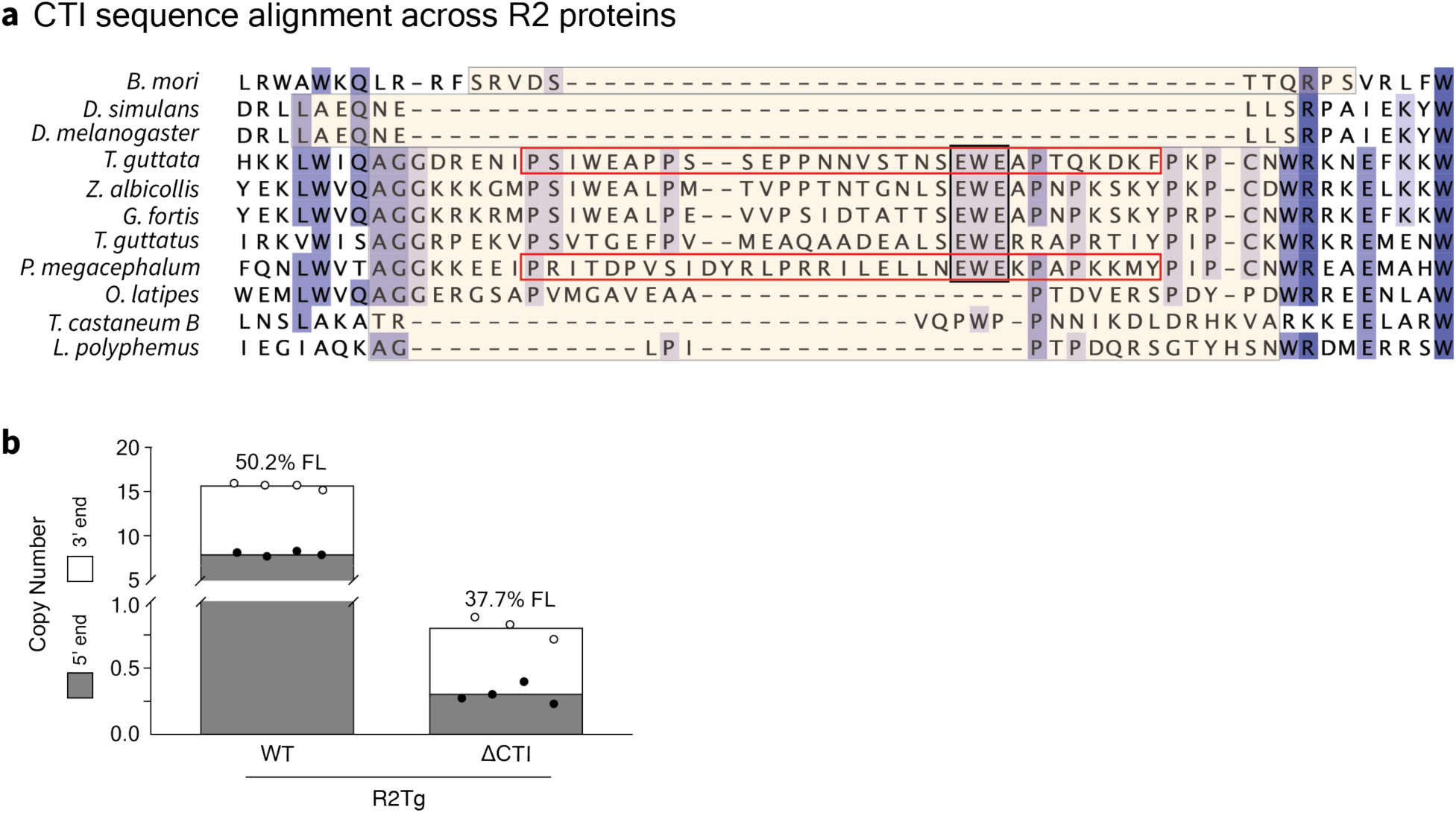
CTI sequence alignment and influence on full-length transgene insertion. (a) The CTI (bounded by peach-colored boxes) and its surrounding sequences were aligned for representative D-clade R2p (rows 1-3) and A-clade R2p (rows 4-11). The CTI boundaries were defined using AlphaFold3 models. The conserved EWE anchor in aligned avian and testudine R2p is highlighted with a black box. Purple shading illustrates relative sequence conservation. Species not given in main text: *Oryzias latipes*, *Limulus polyphemus*, and *Drosophila simulans* or *melongaster* (ref: 13). The red boxes indicate amino acids in R2Pm and R2Tg that were truncated in the ΔCTI mutants. (b) Genomic DNA from cells of Figure 4d, after PRINT with wild-type or ΔCTI R2Tg, was assayed by ddPCR for copy number of the inserted transgene 5′ or 3′ end. Copy numbers are graphed as stacked bars, and the calculated percentage of full-length insertions is indicated above the bars (ref: 17).

**Fig. S8.**
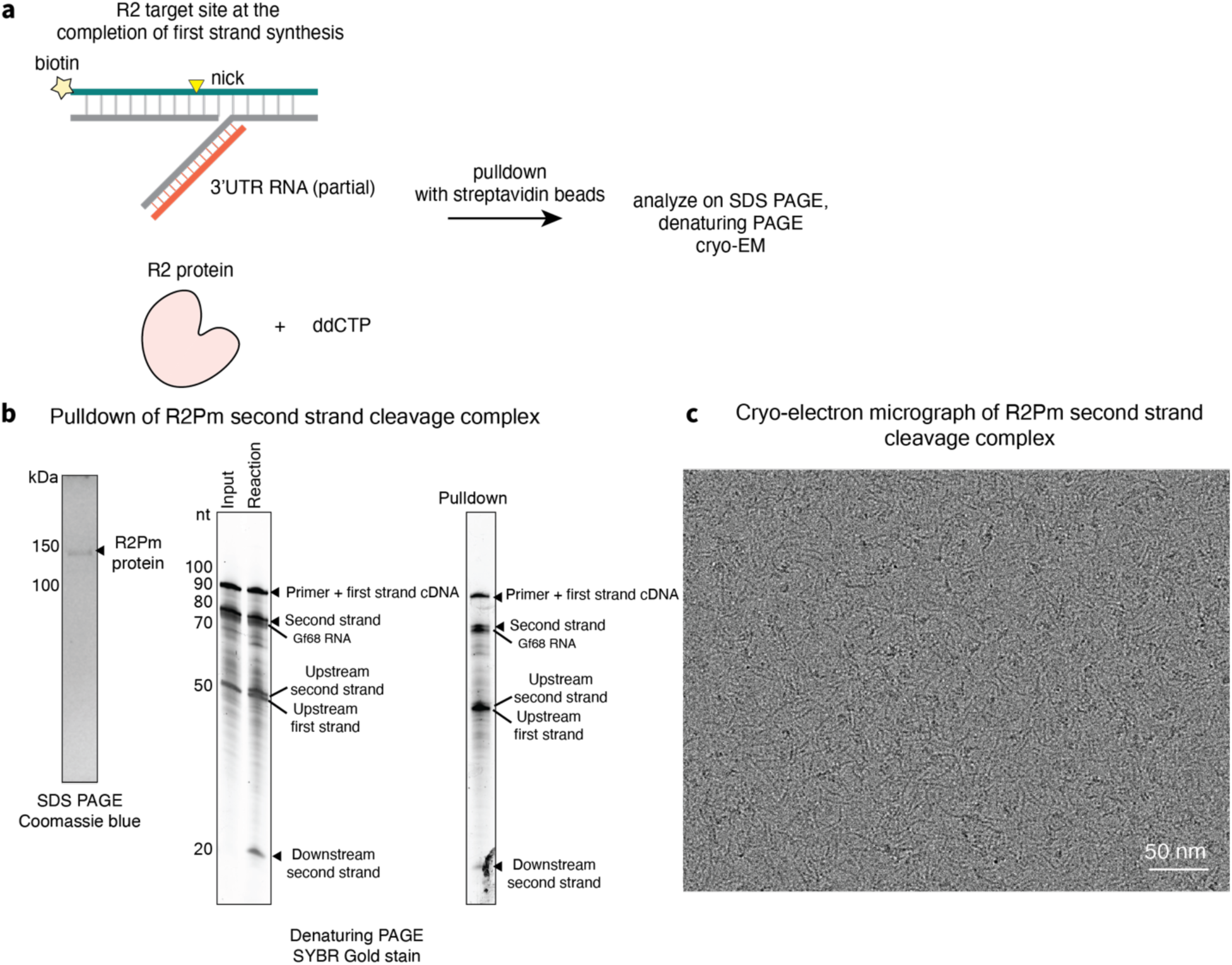
Assembly of second strand nicked complex for cryo-EM analysis. (a) R2Pm was incubated with biotinylated DNA containing the target site and cDNA, with cDNA annealed to template RNA, in a configuration that supports addition of a single ddCTP to complete first strand cDNA synthesis. (b) SDS PAGE protein analysis and denaturing PAGE nucleic acid analysis of the pulldown and elution for the second strand nicked complex with R2Pm. The eluate sample appears to be a mixed population of intact and nicked second strand. (c) Cryo-EM micrographs of the pulldown eluate for R2Pm captured after second strand nicking.

**Fig. S9.**
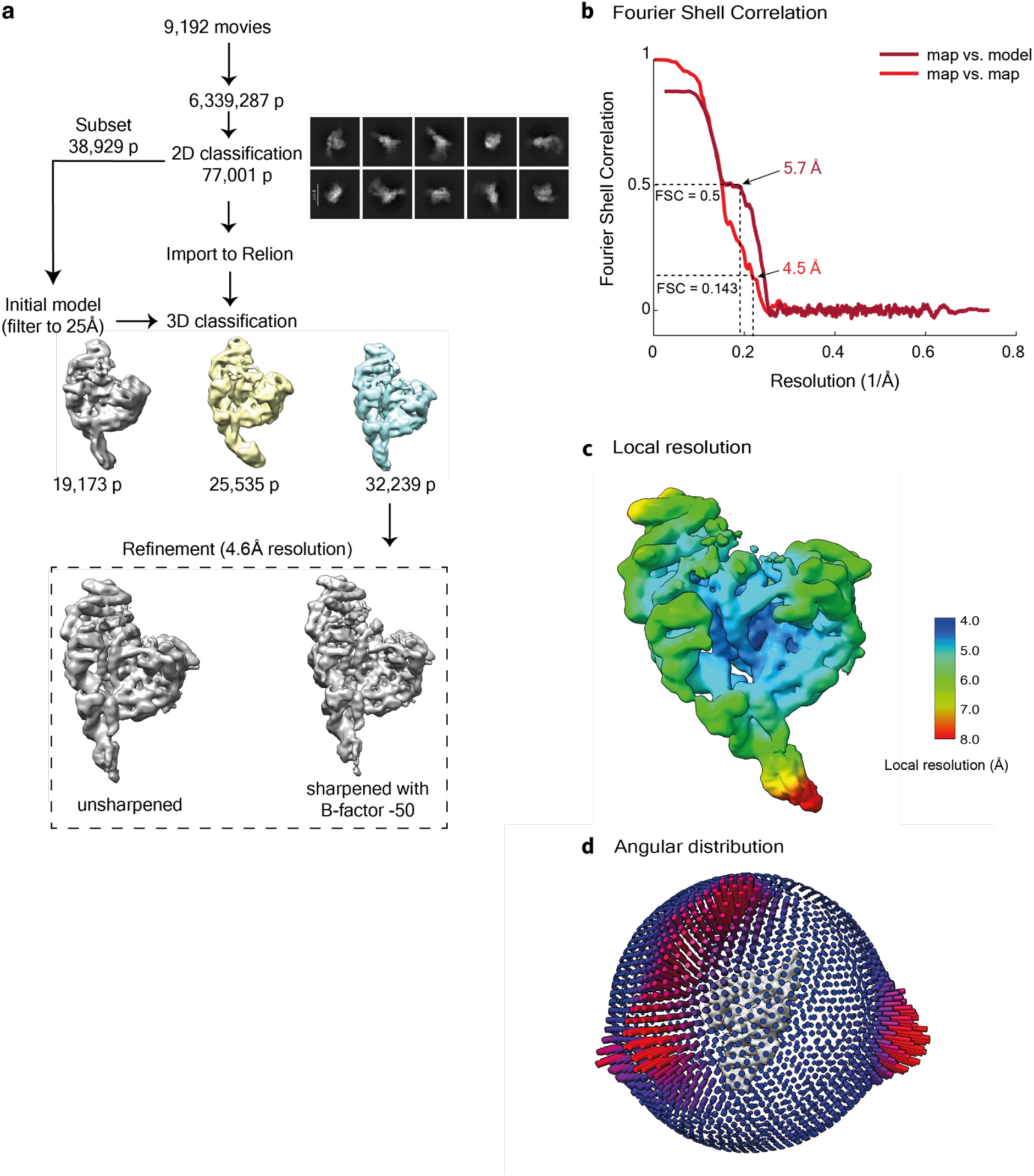
Cryo-EM data processing and resolution estimation for R2Pm second strand nicked complex. (a) Summary of single particle analysis pipeline leading to the reconstruction of the R2Pm second strand nicked complex described in Figure 5. (b) Gold-standard FSC curve and map versus model FSC obtained from the final model after validation in Phenix. (b) Unsharpened density map was colored by local resolution as estimated by Relion 3.1. (c) Particle orientation distribution in the final reconstruction. (c) Particle orientation distribution in the final reconstructions.

